# A protein condensation network contextualises cell fate decisions

**DOI:** 10.1101/2024.04.18.590070

**Authors:** Thomas R Peskett, Sung Sik Lee, Yves Barral

## Abstract

For cells to thrive, they must make appropriate fate decisions based on a myriad of internal and external stimuli. But how do they integrate these different forms of information to contextualise their decisions? Old yeast cells showed an ability to dampen their proliferation as they entered senescence. Conversely, they had an enhanced ability to promote proliferation during escape from pheromone stimulation. A network of nucleoprotein condensation states involving processing bodies (P-bodies) and the prion-like RNA-binding protein, Whi3, controlled these opposing fate decisions. In old but not in young cells, condensation of Whi3 was both necessary and sufficient for senescence entry. In old cells, Whi3 localised to age-dependent P-bodies. Preventing their formation stopped Whi3 condensation from driving senescence entry. Challenging old cells with an external stimulus, pheromone, revealed that the condensates had a second function: potentiating the cell’s ability to trigger escape from the mating pheromone response. These findings identify biomolecular condensation as an integrator of contextual information as cells make decisions, enabling them to navigate overlapping life events.

## Introduction

The decision to start or stop proliferation requires a cell to process a body of information arising from environmental signals and its internal state, which may contradict each other. For example, commitment to proliferation is stimulated by nutrient availability^1–5^, cell size^6^, mitogens^1^ and growth factors, and antagonized by contact inhibition^7^, stressors^8,9^, differentiation signals^10^ and DNA damage^11,12^. How cells prioritize these signals may depend on their own history^13–15^. In metazoans, the decisions that cells ultimately take have profound effects on development and tissue homeostasis. In free living cells, the decisions determine how these organisms adapt to their environments. However, how cells integrate a broad range of signals to decide which fate to engage, remains unclear.

Budding yeast cells (hereafter, yeast) respond to a broad range of pro- and anti-proliferative signals. Depending on nutrient availability, yeast cells engage mitotic proliferation, or enter quiescence^16^ to await the return of better conditions. The proliferative state of haploid yeast cells can itself be repressed by the presence of mating pheromone, which signals the presence of potential mating partners. The cells then make a switch-like decision to arrest their cell cycle in the next G1-phase, and differentiate. They then grow a mating projection, called a shmoo, towards a prospective mating partner^17–19^. Cells that do not find a partner in time are able to stop shmooing and return to the cell cycle^17,20^. Importantly, this time window is sensitive to pheromone concentration and growth conditions^18^.

Overriding these potential decisions, individual yeast cells dampen both their proliferative and mating capacities as they age^21–25^. To divide, yeast cells grow a bud at their surface, the future daughter cell, which is born rejuvenated, while the mother cell retains ageing factors^26^. This replicative ageing limits the mother’s lifespan to 20-30 divisions on average. Before the end of its life, the mother cell enters a senescent state in which it slows down proliferation and ultimately dies^27–29^. The molecular events driving senescence entry and how senescence contributes to cell death are not precisely understood. Senescence might directly reflect damage to the cell cycle machinery or be an adaptive response to ageing. The slow dividing senescent yeast cells are also in a pheromone refractory state. They show a reduced ability to arrest their cell cycle in response to pheromone and exhibit reduced mating abilities^21–23^. How eukaryotic cells in general, and yeast in particular, collect and integrate different information to make context-dependent fate decisions, such as whether to proliferate, mate or enter senescence, is poorly understood.

A large body of work has established that eukaryotic cells take their proliferative decisions at a critical juncture in the G1-phase of the cell cycle called ‘Start’ in yeast and the ‘restriction point’ in metazoans^30^. The molecular networks controlling this point involve the upstream G1 cyclin Cln3 (in yeast) and cyclin D (in metazoans)^30^. In complex with cyclin-dependent kinase (CDK), Cln3/cyclin D activate a transcriptional feedback loop between cyclins, transcriptional repressors, and activators to ensure the irreversibility of cell cycle commitment^31,32^. Various biological contexts interfere with commitment by altering Cln3/cyclin D levels and CDK activity, suggesting that Cln3/cyclin D levels play a key role in contextual decision making^2,3,8,9,11,13,33,34^. Yet, how Cln3/cyclin D expression integrates cellular context information is not fully understood.

The prion-like RNA-binding protein Whiskey 3 (Whi3), a repressor of *CLN3* mRNA translation, is an interesting candidate for linking Cln3 expression to biological context in yeast^35–38^. Indeed, its effects on *CLN3* translation and possibly many other targets affect various fate decisions, including cell cycle entry at a proper cell size, cell cycle arrest in response to pheromone, escape from pheromone arrest, and possibly stress responses^15,35,37,38^.

Whereas Whi3 is diffusely distributed in the cytoplasm of proliferating yeast cells, where it moderately restricts Cln3-CDK activity^38,39^, in stressed cells it localises to P-bodies and stress granules^35,37^. Upon prolonged pheromone exposure, Whi3 forms super-assemblies that release *CLN3* from inhibition, allowing cells to resume proliferation. This super-assembly, called a mnemon, is stable and elicits a self-maintaining pheromone refractory state throughout the subsequent division cycles^15^. Finally, in replicatively old cells Whi3 forms large cytoplasmic puncta that correlate with the reduced sensitivity of these cells to pheromone^23^. Thus, the cellular distribution of Whi3 appears to be sensitive to biological context. Here, we investigate whether and how Whi3’s organisation contributes to context-dependent proliferation decisions.

## Results

### Whi3 promotes senescence entry

To characterise the role of Whi3 in proliferation decisions, we monitored wild type and *whi3Δ* mutant cells (*WHI3* gene deletion) over their entire lifespans using time-lapse microscopy and microfluidics^40^. This setup enabled us to observe individual cells every 15 minutes over 4 days. The wild type cells divided at a regular pace (every hour on average) for the first 75% of their lives, until they entered senescence. Past that point, they drastically slowed their proliferation rate, a hallmark of senescence^27,41–43^ (**Figures 1A and 1B**). Young *whi3Δ* mutant cells initially showed an averaged division time indistinguishable from that of the wild type cells. However, most of the mutant cells failed to slow their proliferation towards the end of life (**Figures 1A and 1B**), suggesting that they failed to enter the senescent state before dying. Classifying senescent cells as those that showed three or more consecutive divisions longer than or equal to 90 minutes each (based on Fehrmann et al.^27^; **Figure S1A**) showed that 92% of wild type cells became senescent before dying (**Figure 1C**). In contrast, only 24% of the *whi3Δ* mutant cells became senescent (**Figure 1C**). Most *whi3Δ* mutant cells ultimately died without slowing their division cycles at all. Importantly, the *whi3Δ* mutant cells that became senescent slowed their division pace as much as wild type cells (**Figure S1A**). Thus, the *whi3Δ* mutation did not alter the division dynamics induced by senescence but impaired senescence entry. Consistent with these data, *whi3Δ* mutant cells were also 28% longer lived than wild type cells (**Figure 1D**). The fact that some *whi3Δ* mutant cells still enter senescence suggests that redundant factors, possibly including Whi3’s paralog, Whi4^44^, compensate for the loss of Whi3 in these cells.

**Figure 1.**
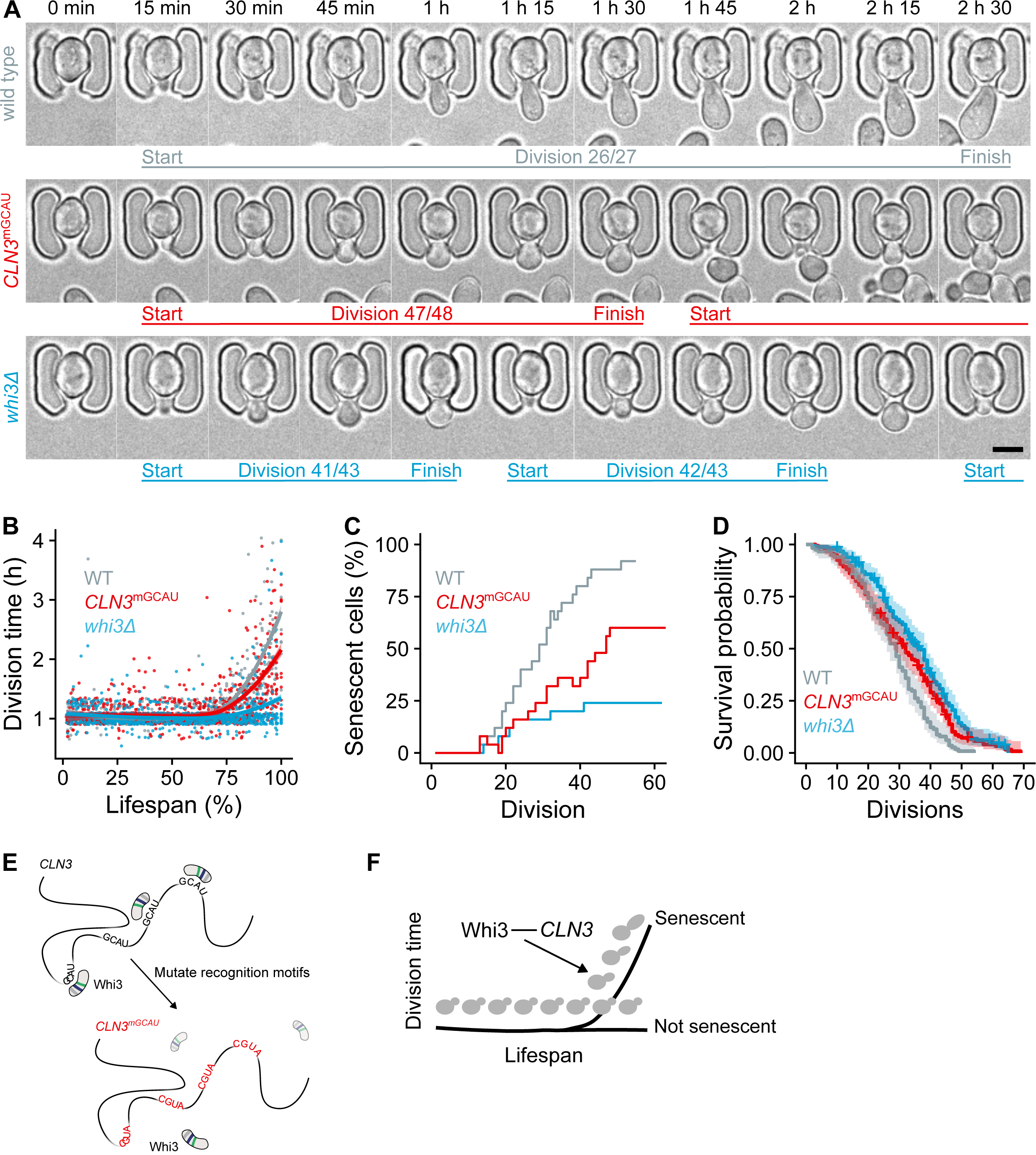
Whi3 promotes senescence entry. (A) Timelapse brightfield microscopy of old single yeast cells undertaking the last divisions of their lives. Comparison between wild type (grey), *CLN3*^mGCAU^ mutant (red), and *whi3Δ* mutant (blue) cells. The start, end and durations of divisions are marked by horizontal lines beneath the images. The division numbers (ages of the cells) are shown above the lines as fractions of the total number of divisions each cell makes (its lifespan). Scale bar, 5 μm. (B) Population averaged division times as a function of percentage lifespan. Smoothed conditional means are shown using the locally estimated scatterplot smoothing method. Shaded areas represent 95% confidence intervals. Data from 823 wild type, 907 *CLN3^mGCAU^,* 973 *whi3Δ* divisions, from 25 wild type, 25 *CLN3^mGCAU^*, and 25 *whi3Δ* cells. (C) Cumulative distribution of senescent cells during replicative ageing. Data associated with (B). Kolmogorov-Smirnov tests: *P* < 1 × 10^−5^ for all comparisons. (D) Kaplan-Meier survival analysis for wild type (grey), *CLN3^mGCAU^* (red), and *whi3Δ* (blue) strains in ageing experiments. Data are the Kaplan-Meier estimate of survival probability (solid lines) and 95% confidence intervals (shaded areas), from 117 wild type, 159 *CLN3^mGCAU^* and 178 *whi3Δ* cells. Log rank tests: wild type vs *CLN3^mGCAU^*, *P* = 0.0011; wild type vs *whi3Δ*, *P* < 0.0001; *CLN3^mGCAU^* vs *whi3Δ*, *P* = 0.047. (E) GCAU to CGUA mutations in *CLN3* disrupt Whi3 binding. (F) Whi3 promotes senescence entry partly through its interaction with *CLN3*. Cells can enter senescence, which involves increasingly prolonged division times, or avoid it, in which case they divide normally until they die.

To better understand how Whi3 promotes senescence entry, we investigated how the expression of its many target transcripts changed with age, compared to the rest of the transcriptome. Using existing ribosome profiling data^45^ and a list of high confidence Whi3 targets^35^ we found that the translation efficiency of Whi3 targets was not significantly different from the rest of the transcriptome in young cells (**Figure S1C**). In old cells, however, Whi3 targets had significantly reduced translation efficiency scores compared to the other transcripts (**Figure S1D**). Notably, many of the targets are involved in cell cycle control^35,38,46^ (**Figures S1B and S1E**) and some of the lowest scores belonged to transcripts of the G1 cyclins Cln2 and Cln3 (**Figure S1E)**. Thus, Whi3 may promote senescence by repressing the translation of its target mRNAs.

We tested this possibility directly by mutating the Whi3 binding sites on the *CLN3* transcript (GCAU motifs, **Figure 1E**). While these mutations leave the Cln3 amino acid sequence intact, they largely prevent Whi3 from repressing the *CLN3* transcript^36^. Remarkably, 40% of the *CLN3^mGCAU^*mutant cells failed to enter senescence before dying, i.e., a 5-fold increase compared to the wild type cohort (**Figures 1A – 1C and S1A**). Some of these cells transiently entered senescence before returning to normal proliferation, indicating that they failed to establish a stable senescent state (**Figure S1A**). Accordingly, the *CLN3^mGCAU^*mutant cells lived 10% longer than wild type cells (**Figure 1D**). Thus, our data indicate that Whi3 promotes senescence entry and does so by repressing the expression of its many targets, the *CLN3* transcript being a prominent one. Supporting this notion, senescent cells slowed down the entire division cycle, which was rescued by the *whi3Δ* and *CLN3^mGCAU^*mutations (**Figure S1F**). This is consistent with Whi3 triggering senescence as a cell state rather than simply slowing progression through G1.

Together these data indicate that senescence entry in old cells is mediated by Whi3 and its ability to repress the expression of its targets, rather than by age-associated damage to the cell cycle engine (**Figure 1F**).

### Whi3’s prion-like domains are required for its effects in old cells

To gain mechanistic insight into Whi3 function, we next dissected the role of its prion-like domains. Together with previous work, our findings indicate that Whi3 functions in both senescence entry and establishing a pheromone refractory state in old cells^23^. Senescence involves the repression of *CLN3* expression whereas the pheromone refractory state requires *CLN3* expression^15^. To rationalise this apparent paradox, we first reasoned that these two functions might involve distinct protein domains. Since the pheromone refractory state involves the two prion-like domains of Whi3, i.e., its disordered glutamine-rich (Q-rich) and asparagine-rich (N-rich) domains^15,47^ (**Figure 2A**), we envisioned that these domains might not be involved in senescence entry. However, when testing this prediction, we found that the *WHI3-ΔQ* mutant cells, which lack the Q-rich domain, showed a reduced ability to enter senescence. Indeed, 39% of these cells failed to enter senescence compared to 14% in the wild type cells (**Figures 2B and S2A**). Full removal of both prion-like domains (*WHI3-ΔQN*) further abrogated cell division slowdown and senescence (**Figures 2B and S2A**). Accordingly, the *WHI3-ΔQ* and *WHI3-ΔQN* mutant cells had longer lifespans than wild type cells (**Figure 2E**), as already reported for the *WHI3-ΔQ* mutation^23^. These effects were not due to the mutant proteins being less stable as their levels remained comparable to that of the full length protein throughout life (**Figure S2B**). However, these mutations substantially impaired the formation of Whi3-3xsfGFP puncta in old cells (**Figures 2C and 2D**), without affecting their occurrence in proliferating young cells (**Figure S2C**).

**Figure 2.**
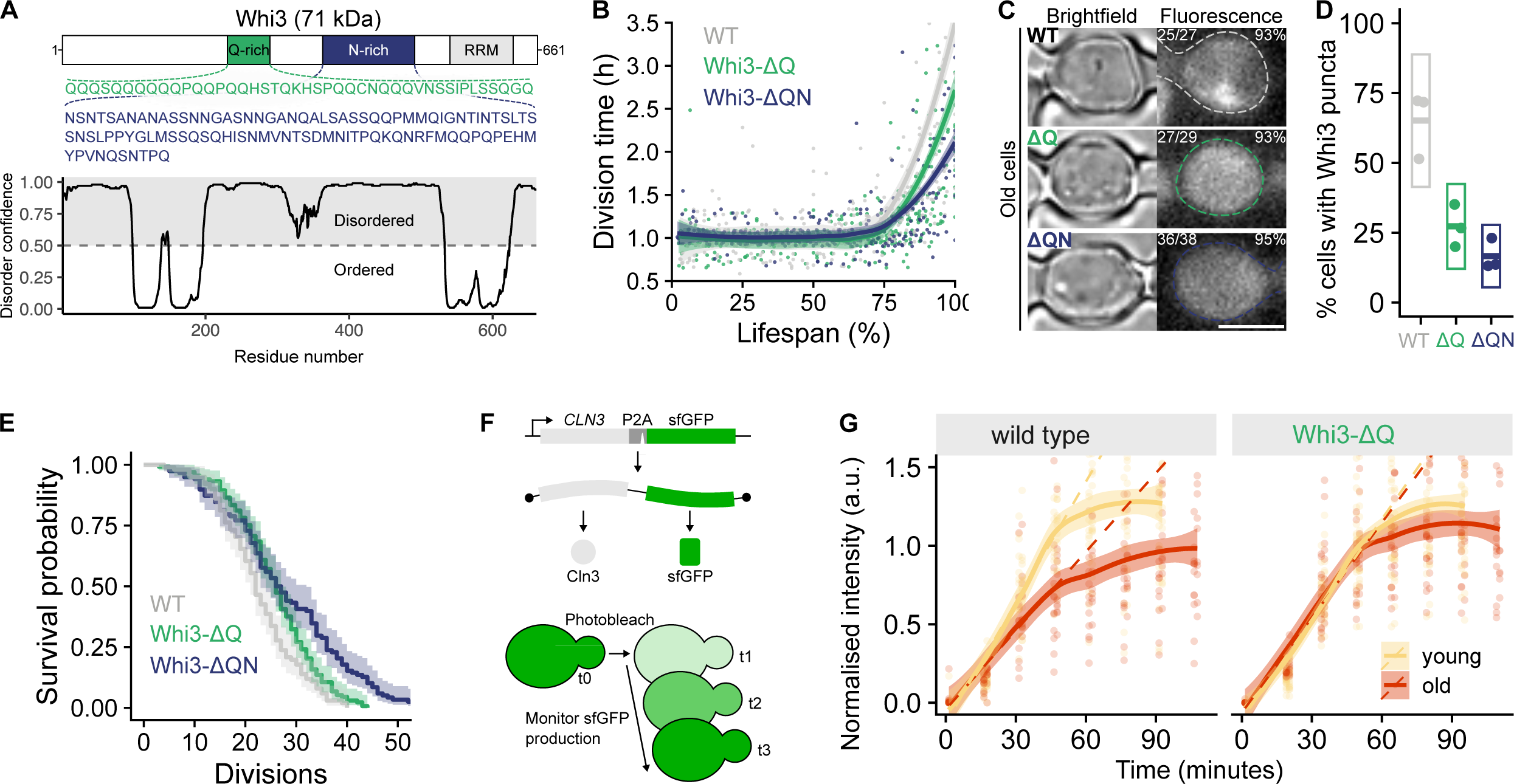
Whi3’s prion-like domains are required for its effects in old cells. (A) Domain organisation of Whi3. Amino acid sequences of the Q-rich (green) and N-rich (purple) domains and location of the RNA recognition motif (RRM, grey). Computational disorder prediction score is aligned to Whi3’s domain organisation. (B) Population averaged division times of the strains shown in (C). Smoothed conditional means are shown using the locally estimated scatterplot smoothing method. Shaded areas represent 95% confidence intervals. Data from 470 Whi3-3sfGFP (WT), 475 Whi3-*Δ*Q-3sfGFP (Whi3-*Δ*Q), 985 Whi3-*Δ*QN-3sfGFP (Whi3-*Δ*QN) divisions, from 21 Whi3-3sfGFP, 18 Whi3-*Δ*Q-3sfGFP, and 28 Whi3-*Δ*QN-3sfGFP cells. (C) Fluorescence microscopy of Whi3-3sfGFP (WT) and Whi3-*Δ*Q-3sfGFP (*Δ*Q) and Whi3-*Δ*QN-3sfGFP (*Δ*QN) in old cells. Ages indicated as fractions and percentages. Note that this strain also contains Edc3-mCherry for reasons that will be explained later in the text. Scale bar, 5 μm. (D) Percentages of cells with Whi3-3sfGFP foci in their last 5 divisions. Points from three independent ageing experiments. Boxes show mean +/-2x standard deviation. Data from 100 Whi3-3sfGFP (WT), 84 Whi3-*Δ*Q-3sfGFP (*Δ*Q) and 75 Whi3-*Δ*QN-3sfGFP (*Δ*QN) cells. Welch’s t-tests: Whi3-3sfGFP vs Whi3-*Δ*Q-3sfGFP, *P* = 0.0140; Whi3-3sfGFP vs Whi3-*Δ*QN-3sfGFP, *P* = 0.0090; Whi3-*Δ*Q-3sfGFP vs Whi3-*Δ*QN-3sfGFP, *P* = 0.128. (E) Kaplan-Meier survival analysis for the strains shown in (B) and (C). Data are the Kaplan-Meier estimate of survival probability (solid lines) and 95% confidence intervals (shaded areas), from 101 Whi3-3sfGFP (WT), 133 Whi3-*Δ*Q-3sfGFP (Whi3-*Δ*Q) and 118 Whi3-*Δ*QN-3sfGFP (Whi3-*Δ*QN) cells. Log rank tests: Whi3-3sfGFP vs Whi3-*Δ*Q-3sfGFP, *P* = 0.0029; WT vs Whi3-*Δ*QN-3sfGFP, *P* < 0.0001; Whi3-*Δ*Q-3sfGFP vs Whi3-*Δ*QN-3sfGFP, *P* = 0.0014. Data associated with (B). (F) Top, a bi-cistronic reporter for monitoring Cln3 production. Bottom, method used to estimate Cln3 production rate. (G) Cln3 production rates in young and old cells of wild type and Whi3-*Δ*Q strains. Smoothed conditional means (solid lines) are shown using the locally estimated scatterplot smoothing method. Shaded areas represent 95% confidence intervals. Dashed lines show a linear model fit to the first three timepoints to estimate the initial rate of production. Individual data points are faded.

Since senescence entry involved repression of *CLN3* function, we next examined whether repression of Cln3 synthesis in old cells depended on Whi3’s prion-like domains. We monitored Cln3 production using a fluorescent reporter^48^ (**Figure 2F**). Consistent with senescence entry being linked to *CLN3* repression, Cln3 production rate declined in old wild type cells (**Figure 2G**). However, while in young *WHI3-ΔQ* mutant cells Cln3’s production rate matched that of young wild type cells (**Figure 2G**), these mutant cells failed to repress Cln3 production in old age (**Figure 2G**). Thus, Cln3 production is indeed repressed in old cells and this requires the Q-rich domain of Whi3.

These findings suggest that Whi3 functions through opposite mechanisms in two cell fate decision processes, senescence entry and mating escape. In the first case it acts by repressing its target genes, particularly *CLN3*, whereas in the second case, it promotes their expression. Paradoxically, both functions depend on the same prion-like domains, and both correlate with Whi3 coalescing into cellular puncta, possibly through condensation.

### Optogenetic Whi3 condensation reconstitutes cell division slowdown and senescence entry

To address this paradox further, we next investigated whether Whi3 condensation into puncta is functionally relevant for senescence entry. Indeed, the puncta formed in old cells might be incidental, and Whi3’s prion-like domains have effects that correlate with, but do not require, their formation. Therefore, we sought to synthetically activate puncta formation by tagging endogenous Whi3 with a Cry2olig tag^49^. Cry2 molecules respond to blue light by increasing their self-association. The resulting increase in valency upon light exposure, can drive tagged proteins containing intrinsically disordered sequences that self-associate to cross a phase boundary and undergo phase separation^50^. This method enabled us to optically induce condensation of Cry2olig-tagged Whi3, called optoWhi3 hereafter (**Figures 3A and S3A**). The light dose used was sufficient to induce optoWhi3 condensation, but not condensation of Cry2olig alone, despite higher expression of Cry2olig compared to optoWhi3 (**Figure S3B**). Blue light induction of optoWhi3 condensates was rapid and reversible, supporting the notion that Whi3 can phase separate *in vivo*^51,52^ (**Figure S3A**). Also, the dynamics of optoWhi3 condensation and dissolution were independent of the number of activation cycles, indicating that these condensates did not mature inside cells (**Figure S3C**).

**Figure 3.**
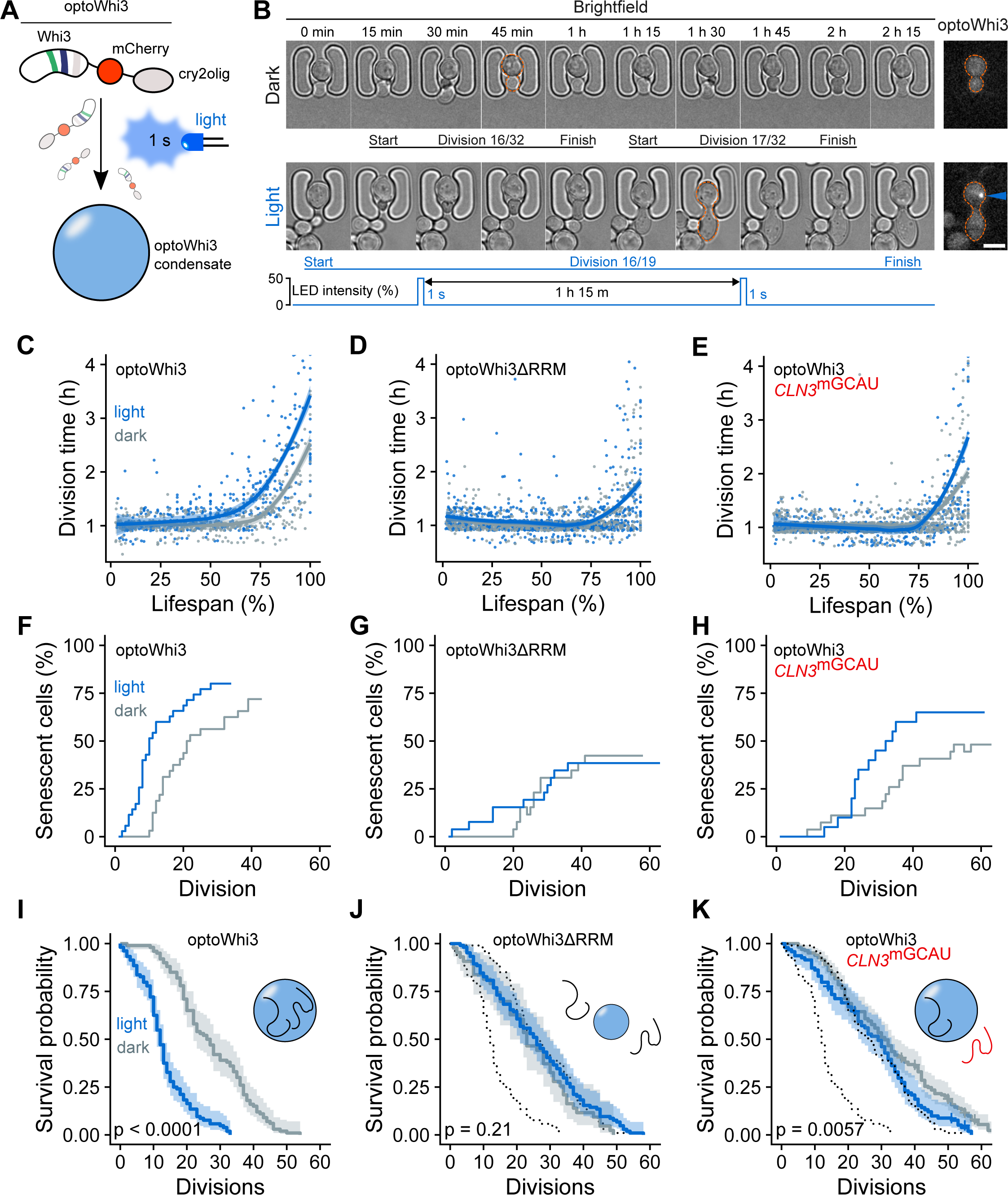
Optogenetic Whi3 condensation reconstitutes slowdown and senescence entry. (A) Light activated condensation of an optogenetic Whi3 construct (optoWhi3). (B) Single cells with identical replicative ages expressing optoWhi3 in either dark (not activated) or light (activated) conditions. The start, end and duration of each division are marked by horizontal lines beneath the images. The division number is shown above the line as a fraction of the total lifespan of the cell. Rightmost panels show optoWhi3 fluorescence signal at the timepoints indicated by an orange dashed outline around the cell. Blue arrow: optoWhi3 condensate. Scale bar, 5 μm. (C) – (E) Population averaged division times of optoWhi3, optoWhi3ΔRRM or optoWhi3 *CLN3^mGCAU^* strains in dark (grey) or light (blue). Smoothed conditional means are shown using the locally estimated scatterplot smoothing method. Shaded areas represent 95% confidence intervals. Data from 511 optoWhi3 (light), 774 optoWhi3 (dark), 875 optoWhi3ΔRRM (light), 778 optoWhi3ΔRRM (dark) 672 optoWhi3 *CLN3^mGCAU^* (light), 1117 optoWhi3 *CLN3^mGCAU^*(dark) divisions, from 35 optoWhi3 (light), 32 optoWhi3 (dark), 26 optoWhi3ΔRRM (light), 26 optoWhi3ΔRRM (dark), 20 optoWhi3 *CLN3^mGCAU^*(light), 27 optoWhi3 *CLN3^mGCAU^* (dark) cells. (F) – (H) Cumulative distribution of senescent cells during replicative ageing for optoWhi3 (F), optoWhi3ΔRRM (G) or optoWhi3 *CLN3^mGCAU^* (H) strains in dark (grey) or light (blue). Data associated with (C) – (E). Kolmogorov-Smirnov tests: optoWhi3, *P* = 0.001; optoWhi3ΔRRM, *P* = 0.002; optoWhi3 *CLN3^mGCAU^, P* < 1 × 10^−7^. (I) – (K) Kaplan-Meier survival analysis for optoWhi3, optoWhi3ΔRRM or optoWhi3 *CLN3^mGCAU^* strains. Data are the Kaplan-Meier estimate of survival probability (solid lines) and 95% confidence intervals (shaded areas). Dotted black lines in (J) and (K) show the superimposed positions of the Kaplan-Meier estimates from (I). *P* values comparing light and dark conditions are from log rank tests. Data associated with (C) – (E). Cartoons depict proposed features of the condensates in each experiment: (I) optoWhi3 condensates bind mRNA; (J) optoWhi3ΔRRM condensates are smaller and do not bind mRNA; (K) optoWhi3 condensates in the *CLN3^mGCAU^* background are normal sized and bind other mRNAs, but not *CLN3* mRNA. Data from 104 optoWhi3 (light), 100 optoWhi3 (dark), 104 optoWhi3ΔRRM (light), 43 optoWhi3ΔRRM (dark), 94 optoWhi3 *CLN3^mGCAU^* (light), 163 optoWhi3 *CLN3^mGCAU^*(dark) cells.

Next, we tested the effects of optoWhi3 condensation by growing colonies in a gradient of blue light (**Figure S3D**). OptoWhi3 condensation did not affect the ability of these young cells to form colonies at any intensity (**Figure S3D**). Next, we induced condensation in cells undergoing ageing in our microfluidic device (**Figure 3B**), exposing them to a low dose of blue light (1 s exposure with a blue LED every 1 h 15 min). Consistent with the colony forming assay, young cells divided indistinguishably whether they were exposed to the light or not (**Figure 3C**). However, as they aged, the exposed cells started slowing their divisions much earlier and more drastically than unexposed ones (**Figures 3B, 3C and S3E**). Moreover, optoWhi3 condensation dramatically shortened the lifespan of the cells (an ∼50% decrease, **Figure 3I**). Cells expressing Cry2olig alone, regardless of light exposure, aged like wild type unexposed cells, demonstrating that neither blue light exposure nor activation of Cry2olig explain the effects (**Figure S3F**). OptoWhi3 condensation did not force a greater percentage of cells into senescence. Rather, it accelerated the senescence entry decision, but only once cells had passed a certain age (**Figure 3F**). Together, these data and the effect of removing the prion-like domains on Whi3 condensation, indicate that in most old cells Whi3 condensation is a determinant of senescence entry.

Consistent with Whi3’s effects being mediated by repression of *CLN3* mRNA and possibly other transcripts, optoWhi3ΔRRM, which lacked the RNA recognition motif, was able to form condensates upon light exposure but did not promote senescence entry or shorten the longevity of the cells (**Figures 3D, 3G and 3J**). Possibly due to the role of RNA in promoting condensation of RNA binding proteins^51,53^, optoWhi3ΔRRM condensed with slower dynamics than optoWhi3 and to a reduced extent (**Figures S3B and S3G**). In the *CLN3*^mGCAU^ mutant cells optoWhi3 condensation was not affected (**Figure S3G**). However, it had a reduced ability to slow down cell division and promote senescence, and little effect on longevity (**Figures 3E, 3H and 3K**). Thus, the ability of Whi3 to bind its RNA targets is essential for Whi3 condensates to promote senescence entry. Furthermore, the enhanced effect of Whi3 condensation in older cells strengthened the idea that some mechanism enables Whi3 condensation to promote senescence entry only in the context of old age.

### Whi3 is enriched in P-bodies

Thus, we sought an age-associated event that could provide the molecular context in which Whi3 condensation engages its effects. We reasoned that this event should satisfy two criteria. First, it should be an ageing-dependent event that occurs late in life, where optoWhi3 condensation has its greatest effect. Second, it should be required for senescence entry. A recent preprint shows that old yeast cells accumulate P-bodies late in life^54^. P-bodies are cytoplasmic biomolecular condensates that contain translationally repressed RNAs and proteins involved in RNA processing^55^. In *S. cerevisiae*, they form in response to stressors such as glucose starvation and heat shock. Furthermore, Whi3 localises to P-bodies in those situations^35,37^. Therefore, we asked whether P-bodies matched the criteria set out above.

In agreement with P-bodies fulfilling the criteria, fluorescence microscopy confirmed that Edc3-GFP (a P-body marker), condensed in old cells, whereas Pub1-GFP (a stress granule marker) did not (**Figure S4A**). P-bodies generally formed in the last quarter of a cell’s life when the cell entered senescence (**Figures S4B and S4C**). Indeed, there was a positive correlation between P-body condensation and the division time of the cell (Spearman’s rank correlation rho = 0.77, p = 2.2 × 10-16, **Figure S4D**). This correlation was stronger than that between a cell’s age and its division time (Spearman’s rank correlation rho = 0.44, p = 3.4 × 10-11). These data show that P-body formation occurs late in life and correlates with slowing cell divisions.

P-bodies could be a cause or a consequence of slowing cell divisions. In support of the former, P-body inheritance appeared to be a strong determinant of senescence entry. Whereas P-bodies were generally retained in the ageing mother cell over its lifespan (**Figures S4E – S4G**), some mother cells occasionally lost their P-bodies to their daughter cells during division. In these cases, the mother cells had rejuvenated division times (**Figure S4H**), consistent with Choi et al.^54^. This might explain the ability of some senescent cells to recover the ability to divide normally^28^. These observations indicate that P-bodies themselves, rather than related diffusible structures, help establish the senescent state.

We therefore investigated whether Whi3 and P-bodies are functionally related in old cells. Using fluorescence microscopy, we found that Whi3 puncta in old cells largely overlapped with the age-induced P-bodies, marked by Edc3-mCherry (**Figures 4A, 4B and S5A**). Whi3’s prion-like domains were required for this colocalisation because it was largely lost in *WHI3-ΔQ* and *WHI3-ΔQN* mutant cells (**Figures 4A and 4B**). Importantly, P-body condensation was not significantly affected in these mutant cells, establishing that Whi3 colocalisation with P-bodies is not required for their formation (**Figures 4A and 4C**). Consistent with this, Whi3 enrichment did not correlate with Edc3-mCherry intensity, suggesting that Whi3 does not cooperatively assemble with core P-body components^56^ (**Figure S5B**). Overall, these data indicated that P-bodies might provide the molecular context required for Whi3 to trigger senescence entry specifically in old cells.

**Figure 4.**
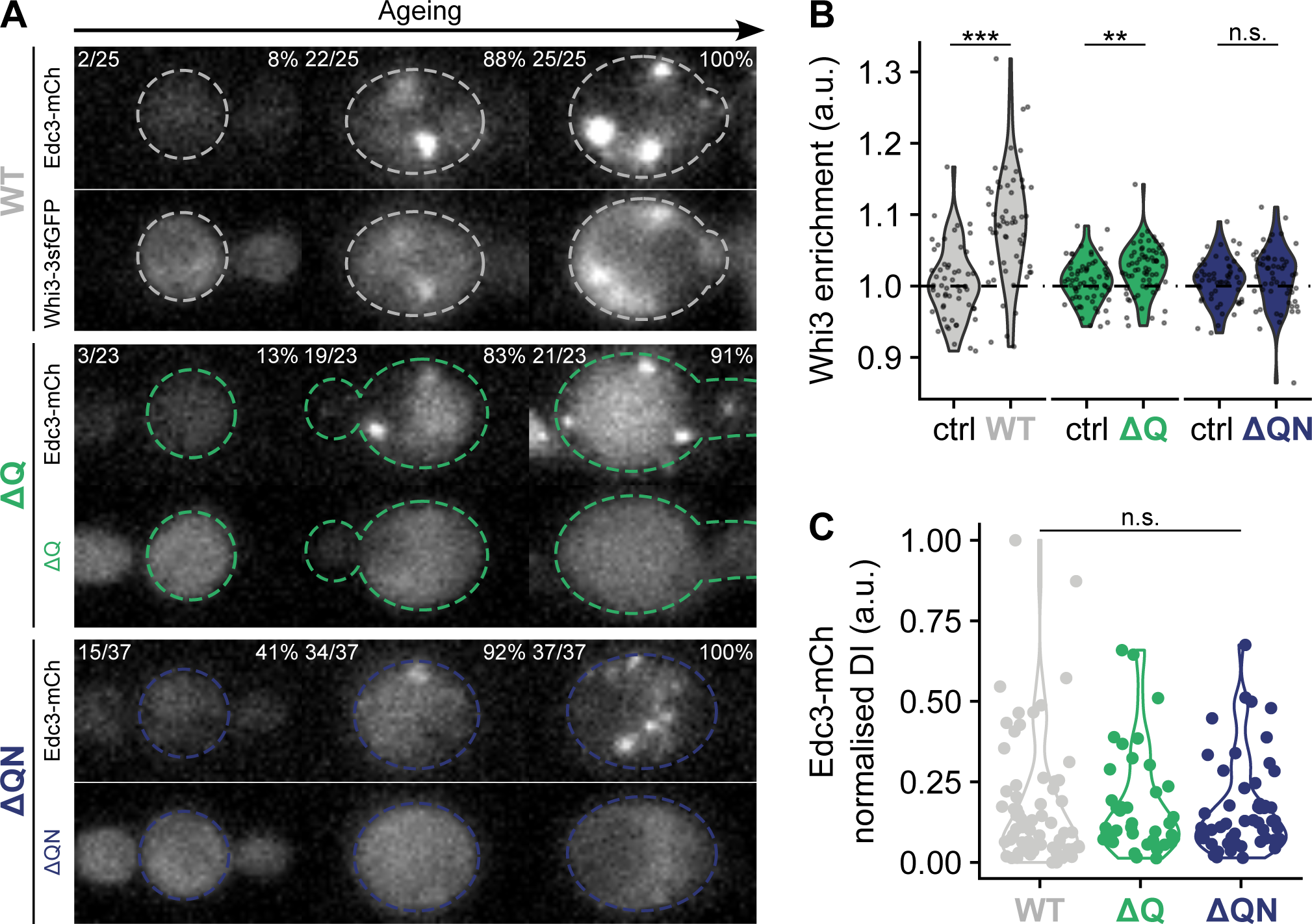
Whi3 is enriched in P-bodies. (A) Timelapse fluorescence microscopy of individual cells expressing Edc3-mCherry and Whi3-3sfGFP and the Whi3-ΔQ-3sfGFP (ΔQ) and Whi3-ΔQN-3sfGFP (ΔQN) mutant versions. Cell ages are shown as fractions and percentages of lifespans. Scale bar, 5 μm. (B) Quantification of Whi3 enrichment in P-bodies. Data related to (A). Data points are single cells in their last 5 divisions. Ctrl: control, enrichment score from two randomly selected areas of the cytoplasm. Welch’s t-test. *** *P* < 0.001, ** *P* < 0.01, n.s. *P* > 0.05. (C) P-body condensation quantified by measuring Edc3-mCherry dispersion index (DI). Data points are single cells in their last 5 divisions. Welch’s t-test. n.s. *P* > 0.05.

### P-bodies promote cell division slowdown and senescence entry

To test this idea, we investigated whether P-body formation is required for old cells to enter senescence. The P-body component Lsm4 (homolog of human LSM4), a subunit of the Lsm1-7 complex involved in mRNA decapping^57,58^, harbours a prion-like tail at its C-terminus. Similarly, Edc3 (homolog of human EDC3), a decapping activator, has a C-terminal dimerisation ‘YjeF’ domain (structurally related to the N-terminal domain of the bacterial YjeF protein). In young cells, P-body formation can be disrupted by truncating the C-termini of Lsm4 (Lsm4ΔC) or Edc3 (Edc3ΔYjeF)^59^ (**Figure 5A**). These mutations have mild effects on the core decapping functions of these proteins, and mainly affect the multivalent interactions required for P-body formation^59–63^. Thus, we asked whether these mutations or combinations of them could also disrupt age-induced P-bodies. Both *LSM4ΔC* and *EDC3ΔYjeF* mutant cells formed fewer P-bodies as they aged and the *LSM4ΔC EDC3ΔYjeF* double mutant cells hardly formed visible P-bodies, showing that the mutations additively disrupt age-induced P-body condensation (**Figures 5B and 5C**).

**Figure 5.**
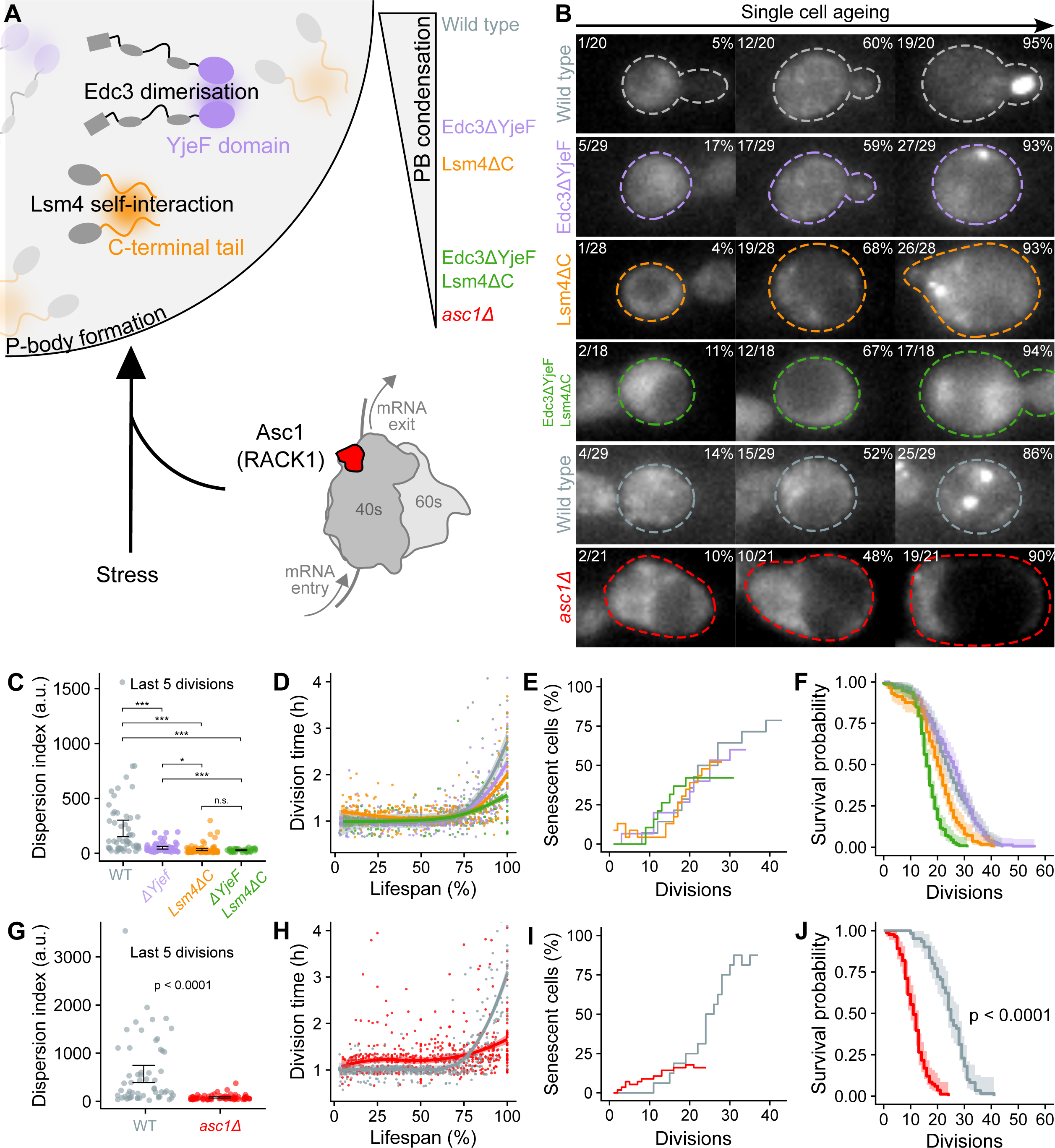
P-bodies promote cell division slowdown and senescence entry. (A) P-body formation is supported by Edc3 dimerisation through its C-terminal YjeF domain, and Lsm4 self-interaction through its C-terminal prion-like tail. Asc1/RACK1, supports P-body formation under DNA replication stress and perhaps other processes. Edc3ΔYjeF and Lsm4ΔC protein truncations partially disrupt P-body formation in old cells. Combining both almost completely disrupts P-body formation, as does an *asc1Δ* mutation, giving a set of mutant cells with graded P-body disruption. (B) Timelapse fluorescence microscopy of ageing single cells expressing Dcp2-GFP, which marks P-bodies. Cell ages are shown as fractions and percentages of their total lifespan in the top left and top right corners of each panel, respectively. Scale bar, 5 μm. (C) P-body condensation of cells in their last 5 divisions quantified by measuring Dcp2-GFP dispersion index. Whiskers show 95% confidence intervals about the mean. **** *P* < 0.0001, *** *P* < 0.001, * *P* < 0.05, n.s. *P* > 0.05. (D) Population averaged division times of the strains shown in (C). Smoothed conditional means are shown using the locally estimated scatterplot smoothing method. Shaded areas represent 95% confidence intervals. Data from 330 wild type, 292 Edc3ΔYjeF, 433 Lsm4*Δ*C, 287 Edc3ΔYjeF Lsm4*Δ*C divisions, from 14 wild type, 15 Edc3ΔYjeF, 23 Lsm4*Δ*C, 19 Edc3ΔYjeF Lsm4*Δ*C cells. (E) Cumulative distribution of senescent cells during replicative ageing. Data associated with (D). Kolmogorov-Smirnov tests: wild type vs Edc3ΔYjeF, *P* = 0.002; wild type vs Lsm4*Δ*C, *P* = 3 × 10^−5^; wild type vs Edc3ΔYjeF Lsm4*Δ*C, *P* = 2 × 10^−5^; Lsm4*Δ*C vs Edc3ΔYjeF *P* = 0.018; Lsm4*Δ*C vs Edc3ΔYjeF Lsm4*Δ*C *P* = 0.0059; Edc3ΔYjeF vs Edc3ΔYjeF Lsm4*Δ*C *P* = 0.059. (F) Kaplan-Meier survival analysis for strains shown in (C) and (D). Data are the Kaplan-Meier estimate of survival probability (solid lines) and 95% confidence intervals (shaded areas), from 131 wild type, 141 Edc3ΔYjeF, 112 Lsm4*Δ*C, and 112 Edc3ΔYjeF Lsm4*Δ*C cells. Log rank tests: wild type vs Edc3ΔYjeF, *P* = 0.63; all other pairwise comparisons between strains have *P* < 0.0001. (G) P-body condensation for cells in their last 5 divisions quantified by measuring Dcp2-GFP dispersion index. Whiskers show 95% confidence intervals about the mean. (H) Population averaged division times of the strains shown in (G). Smoothed conditional means are shown using the locally estimated scatterplot smoothing method. Shaded areas represent 95% confidence intervals. Data from 417 wild type and 641 *asc1Δ* divisions, from 16 wild type and 56 *asc1Δ* cells. (I) Cumulative distribution of senescent cells during replicative ageing. Data associated with (H). Kolmogorov-Smirnov test: *P* = 1 × 10^−5^. (J) Kaplan-Meier survival analysis for strains shown in (G) and (H). Data are the Kaplan-Meier estimate of survival probability (solid lines) and 95% confidence intervals (shaded areas), from 61 wild type and 83 *asc1Δ* cells. *P* value of a log rank test is shown.

We used these mutants to ask whether P-body assembly is required for senescence entry. Like the *whi3Δ*, *WHI3-ΔQ* and *WHI3-ΔQN* single mutant cells, the P-body disruption mutant cells did not slow their divisions as dramatically as wild type cells in old age, and fewer cells entered senescence (**Figures 5D and 5E**). In contrast to Whi3 mutants, P-body disruption either shortened or did not dramatically affect lifespan (**Figure 5F**). Likewise, cells lacking the Asc1 protein (homolog of human RACK1), a ribosomal protein involved in P-body formation under conditions such as DNA replication stress^64^ (**Figure 5A**) did not form P-bodies with age (**Figures 5B and 5G**). These cells largely failed to slow down cell division and enter senescence prior to death (**Figures 5H and 5I)**. They also had a reduced lifespan (**Figure 5J**). Overall, these data reveal that P-body formation extends the longevity of ageing cells on one hand and promotes their transition to senescence on the other (**Figure S5C and S5D**). Thus, P-bodies fulfilled the criteria for an age-associated event that could enable Whi3 condensation to exert its effects.

### P-bodies empower Whi3 condensation to slow proliferation in old cells

To test whether P-bodies indeed provide the molecular context for Whi3-driven senescence entry, we introduced P-body disruption mutations in the optoWhi3 strain. Importantly, disrupting P-body formation did not affect optoWhi3 condensation (**Figure S6A**). In contrast to cells with a wild type background, optoWhi3 condensation had no effect on cell cycle progression in cells with near complete P-body disruption (*LSM4ΔC EDC3ΔYjeF* and *asc1Δ*), as determined by measuring division times (**Figures 6A, S6B, S6E and S6F**). In the *LSM4ΔC* and *EDC3ΔYjeF* single mutant cells (partial P-body disruption), the effects of the optoWhi3 condensates on cell cycle progression were partially diminished (**Figures 6A, S6C and S6D**). With partial P-body disruption, optoWhi3 condensation was sufficient to drive more cells into senescence, but this ability was lost upon full P-body disruption (**Figures 6B and S6G – S6K**). Consistent with these effects, optoWhi3 condensation did not shorten the lifespans of mutant cells with full P-body disruption and did so only to a limited extent in those with partial P-body disruption (**Figures 6C and S6L – S6P**). Thus, below some threshold of P-body formation, Whi3 condensation does not promote senescence entry. These effects establish a cooperative process between Whi3 and P-body components that relies on protein condensation driven by multivalency.

**Figure 6.**
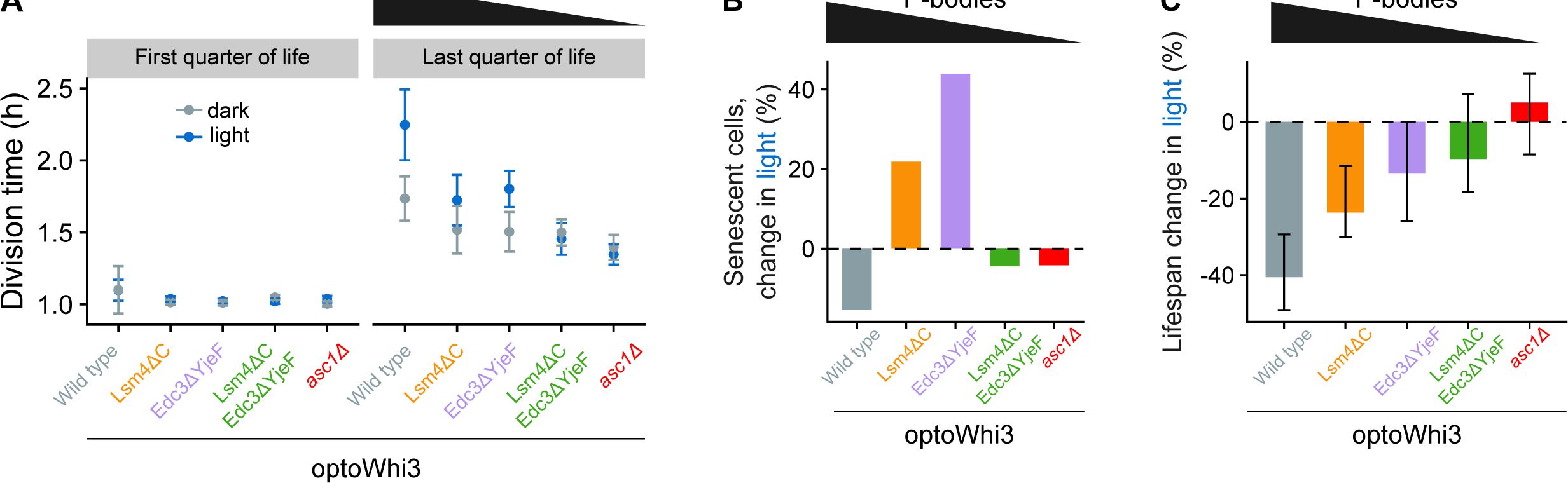
P-bodies empower Whi3 condensation to slow proliferation of old cells. (A) Averaged division times in the first and last quarter of a cell’s life in light (optoWhi3 condensation activated) and dark (optoWhi3 condensation not activated) conditions, with different levels of P-body disruption. Data are mean division times (points) and 95% confidence intervals (whiskers), related to Figures S6B – S6F. (B) How optoWhi3 condensation changes the percentage of cells that enter senescence in blue light compared to dark, in genetic backgrounds with different levels of P-body disruption. Dashed line indicates zero. Related to Figures S6G – S6K. (C) Summary of percentage lifespan reduction caused by optoWhi3 condensation in strains with different levels of P-body disruption. Data are percentage differences in median lifespans between dark and light conditions, from Kaplan-Meier survival analyses, related to Figures S6L – S6P. Whiskers indicate 95% confidence intervals after error propagation. Dashed line indicates insufficient evidence for lifespan reduction if a confidence interval overlaps.

### P-bodies promote a mating escape decision which depends on the age of the cell

Together, our data indicate that in old cells, cooperation between Whi3 condensation and P-bodies promotes senescence entry by repressing expression of Whi3 target transcripts, such as *CLN3* mRNA. The same cells have been found by us and others to be largely sterile^21–23^. This is paradoxical because *CLN3* repression should facilitate the pheromone response^15,65^. Furthermore, the pheromone refractory state requires Whi3’s super-assembly, which promotes *CLN3* expression^15^. This initially appeared incompatible with the P-body-associated pro-senescence state reported here.

To rationalize this apparent paradox, we reexplored the pheromone response behaviour of old cells. We permanently switched old cells to pheromone-containing medium and observed their responses at high temporal resolution (**Figure 7A**). Unexpectedly, the old cells did respond to pheromone. However, they rapidly escaped the pheromone response, much faster than young cells (**Figures 7A and 7B)**. The older the cells, the more rapidly they escaped and entered the pheromone refractory state (**Figure 7B**). Thus, most old cells were not in the pheromone refractory state prior to pheromone exposure. Instead, they were primed to enter it quickly in response to pheromone, and hence, did so faster than young cells. We conclude that, like the decision to enter senescence, the decision to escape the pheromone response depends on the age context of the cell. Classical experimental approaches relying on purified old cells and microdissection may have missed this rapid escape phenotype^21–23^. These results, and Whi3’s function in senescence entry, indicate that ageing alone does not induce Whi3 super-assemblies.

**Figure 7.**
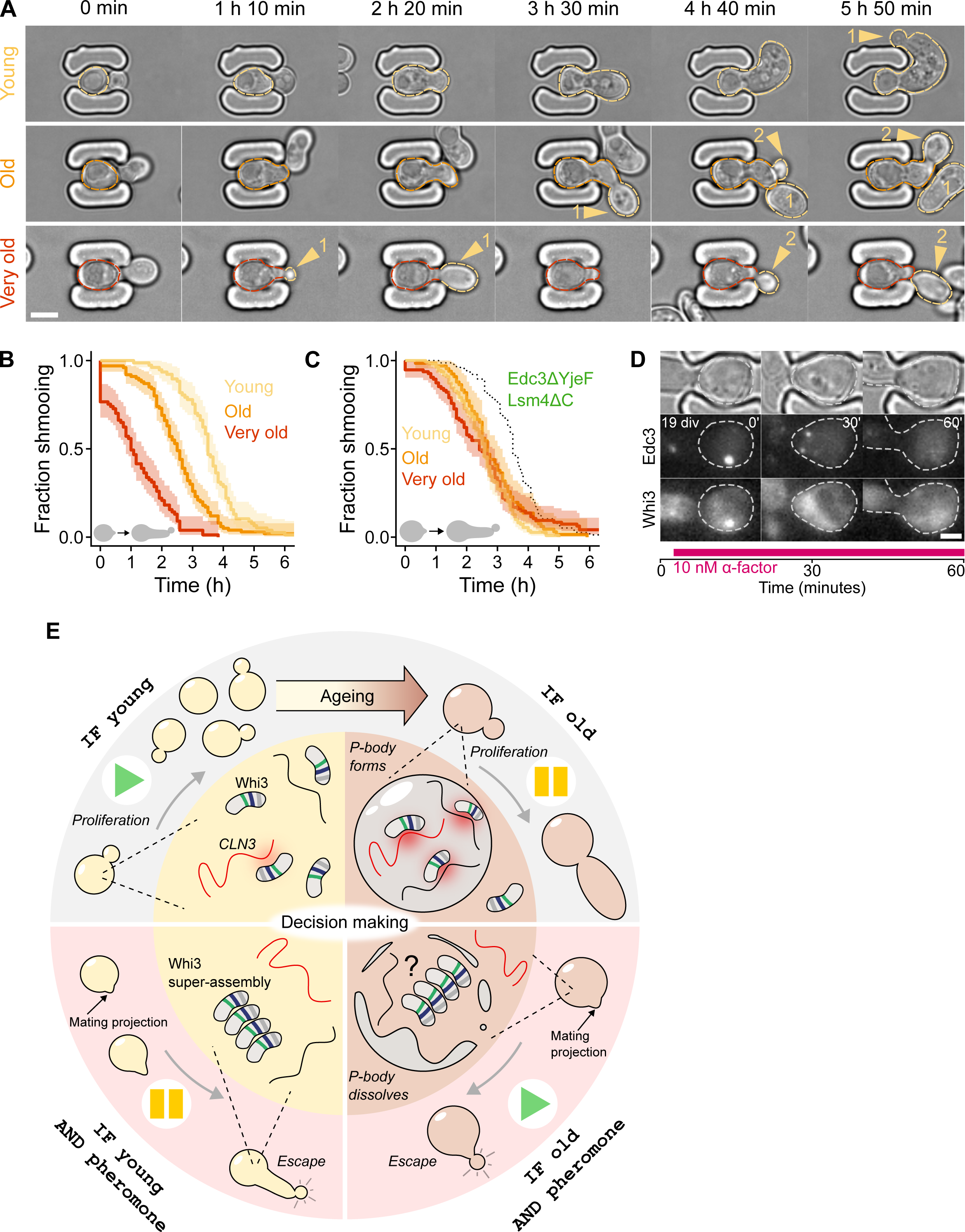
P-bodies promote a mating escape decision which depends on the age of the cell, and a model for contextual decision making. (A) Timelapse brightfield microscopy of young, old, and very old cells responding to pheromone, growing mating projections, and escaping the response. New daughter cells (yellow arrows) mark the escape from mating and return to proliferation. Scale bar, 5 μm. (B) Pheromone response and escape dynamics of cells with different ages. Data are the Kaplan-Meier estimates of escape probabilities (solid lines) and 95% confidence intervals (shaded areas), from 77 young, 100 old and 77 very old cells. (C) Pheromone response and escape dynamics of Edc3ΔYjeF Lsm4*Δ*C mutant cells with different ages. Dotted black trace shows the position of the curve for young wild type cells, in (B). Data are the Kaplan-Meier estimates of escape probabilities (solid lines) and 95% confidence intervals (shaded areas), from 110 young, 66 old and 95 very old cells. (D) Timelapse microscopy of P-body dissolution after pheromone exposure. Cells were switched to new medium containing 10 nM pheromone (⍺-factor), in parallel with a control sample that was switched to new medium without pheromone (Figure S7B). Fluorescence images are maximum intensity projections of Edc3-mCherry and Whi3-3sfGFP across four slices spanning a depth of 2 μm. ’div’ indicates the age of the cell at the media switch. Numbers with inverted commas are minutes after media switch. Scale bar, 2.5 μm. (E) Model of the protein condensation network that informs cell fate decisions. If cells are young and there is no pheromone in their environment, they proliferate rapidly. Whi3 is in an uncondensed state, and it inhibits *CLN3* at a basal level. As cells age, P-bodies containing Whi3 form, which enhances *CLN3* inhibition (and possibly other RNAs). This slows proliferation and promotes senescence entry. If pheromone in the cell’s environment indicates a mating partner is nearby, young cells initiate a prolonged mating attempt that ends due to Whi3 super-assembly and *CLN3* release^15^. In contrast, old cells initially respond to pheromone, but age-induced P-bodies dissolve and promote rapid escape from mating. Question mark: do P-bodies prime Whi3 for conversion to its super-assembled state?

To better understand what primes old cells to rapidly escape the pheromone response, we tested whether P-bodies influence the escape decision. We compared the pheromone responses of wild type cells with *LSM4ΔC EDC3ΔYjeF* double mutant cells, as these do not form P-bodies in old age. Young mutant cells escaped slightly faster than young wild type cells, suggesting that P-body formation might initially help maintain the G1 arrest in young cells (**Figure 7C**). Moreover, the mutant cells escaped at the same rate independent of their ages, such that old cells escaped as slowly as young cells (**Figure 7C**). Thus, the P-bodies of old cells promote the mating escape decision, in addition to the senescence entry decision. Strikingly, the P-bodies and condensed Whi3 rapidly dissolved upon pheromone treatment (**Figures 7D, S7A and S7B**).

Consistent with pheromone treatment leading to P-body dissolution, when wild type cells were treated with a 1 h 50 min pulse of pheromone (20 nM, ensuring that all cells responded and escaped regardless of age) the survival probability of very old cells was unchanged, but their cell division slowdown was less pronounced (**Figures S7C and S7D**). Furthermore, this treatment temporarily protected young and moderately old cells from death, causing their survival probability to plateau for about 6 divisions, before falling again (**Figures S7E and S7F**). Their average lifespans were unaffected, although we did not observe the cells reaching extremely old ages, possibly due to their decreased retention in the microfluidic device as they grew larger during shmooing. This suggests that escape from pheromone-induced arrest alleviates ageing effects but does not alter the underlying ageing process, analogous to a health span increase.

## Discussion

How do cells sense their internal state and integrate that with other information from their environment, to decide between different fates, such as proliferation, mating and senescence? We propose that Whi3 and P-bodies are components of a protein condensation network that integrates information about a cell’s internal state and its surroundings, allowing it to make context-dependent decisions. This network converges on specific mRNAs to control their expression, which determines the final decision (**Figure 7E**). We suggest that Whi3 condensation inhibits *CLN3* and possibly other mRNAs. This effect is strongly enhanced by Whi3’s association with P-bodies, at least in old cells. P-bodies might enhance Whi3’s activity, bring it closer to its target mRNAs, and/or exclude mRNA-ribosome interactions. Studies linking P-bodies to translation repression are consistent with this idea^66–69^, and our genetic experiments suggest that Whi3 needs to bind RNA for its condensation to affect cell division (**Figures 1 and 3**). Ultimately, these processes slow cell proliferation and promote senescence entry.

The same network also contextualises mating decisions. In young cells which do not contain P-bodies, the pheromone response causes Whi3 to super-assemble slowly^15^. Eventually the cells escape mating and return to proliferation due to the release of *CLN3* from inhibition^15^. In contrast, in old cells, Whi3-enriched P-bodies speed up the escape decision. One possibility is that P-bodies prime Whi3 for super-assembly via a liquid-solid phase transition, which would be consistent with P-bodies having liquid-like properties^70^, Whi3’s Q-rich domain potentiating escape in old cells and forming amyloid fibres *in vitro*^23,47^, and liquid-to-solid conversion of *Ashbya gossypii* Whi3 condensates *in vitro*^51^. This model would tie different decisions to the material properties of the network components, a strategy that could have evolved in other organisms^71^. P-body dissolution or an inability to form them could prevent senescence decisions and explain why some young and middle-aged cells gain a temporary survival advantage after pheromone exposure (**Figures S7E and S7F**). Aside from the underlying mechanism, priming of Whi3 super-assemblies by P-bodies would explain how Whi3’s Q-rich domain can inhibit proliferation in old cells that are cycling, but promote proliferation in old cells that are exposed to pheromone. Our findings add to a growing list of complex unicellular behaviours that are normally associated with networks of cells performing cognitive tasks: sensing, decision making, learning and memory^14,15,72–75^.

Many changes in a cell’s environment or within the cell itself cause proteins to accumulate in condensates. Recent breakthroughs have highlighted some of their biological functions^76–83^. Yet, some of the most ubiquitous condensates, such as P-bodies and stress granules, have poorly defined functions. Our findings suggest that P-bodies may be used by a cell to probe its internal state during decision making. P-bodies are ubiquitous in nature and their conserved regulation^84^ indicates that our findings might also have implications in more complex organisms.

P-bodies often form or dissolve during important developmental transitions^85^. Their dissolution during yeast mating (**Figure 7D**) resembles the dissolution of similar condensates when arrested oocytes are fertilised^86,87^. Recently, they have been linked to meiotic exit in *Arabidopsis*, embryonic stem cell potency in mouse, and the adaptability of cancer cells^66,88,89^. Their functions are still debated^90^, but a common theme that emerges from these and our observations is decision making.

Biological condensates have several features that could directly facilitate decision making. Phase separation is a driving force behind many condensates including P-bodies^91^. Crossing the phase boundary between different phases involves a sharp, decision-like, thermodynamic transition^18^, which could allow cells to make robust decisions in noisy environments^92^. This behaviour is exemplified by proteins like Pab1 and Ded1 which rapidly condense upon heat shock, acting as binary stress sensors^93,94^. Furthermore, condensates can directly respond to physical conditions such as osmolarity, temperature and acidity, and biological signals such as nucleic acid concentrations and post-translational modifications^95^. The combination of these signals, arising from the cell’s context, could steer the cell through a phase diagram, allowing it to integrate multiple sources of information to make decisions. The context-dependent compositions of both P-bodies and stress granules support this idea^96–98^. An important addition of our model is that the Whi3/P-body condensation network injects contextual information into multiple decisions to achieve different outcomes. It thereby enables cells to carry out logical operations, combining operators such as IF and AND (**Figure 7E**).

Decision mechanisms based on protein condensation might also come at a cost. Many age-associated disease proteins convert from liquid-like condensates to solid-like aggregates *in vitro*^99–102^. Thus, condensates intended for making age-dependent decisions might nucleate aggregation of these mutant proteins or have their material properties and functions affected by them^103^. Indeed, recent work shows that alpha-synuclein toxicity, involved in Parkinson’s disease, is linked to P-bodies^104^. Understanding the context-dependent functions of condensates should help clarify their links to disease.

Our findings suggest that two archetypal yeast ageing phenotypes – senescence and sterility – arise from decisions made by the cell. In support of this idea, many cells avoid senescence entry if components of the decision network (Whi3, *CLN3*, and P-bodies) are perturbed. Furthermore, when mother cells lose age-induced P-bodies during cell division, their next divisions are revived^54^ (**Figure S4H**). In other words, senescence is likely an adaptive response rather than irreversible loss of a cell’s division potential. Similarly, old cells do not lose their ability to mate, instead they escape mating more quickly than young cells. Recent work shows that yeast ageing trajectories bifurcate early between two distinct modes of ageing that involve mitochondrial dysfunction or chromatin instability^105^. Together with senescence, which applies to both modes of ageing^105^, and mating decisions, hallmarks of ageing yeast cells appear to result from a complex landscape of decisions rather than general deterioration. The situation might be similar in metazoans^106,107^. An important next step is to understand the causes, consequences, and benefits of these decisions for cells.

We hypothesise that P-bodies contextualise many decisions, not just those associated with ageing. Until now, work has reasonably focused on one type of P-body inducing stimulus at a time. Future work could explore the effects of multiple overlapping stimuli, where contextual decision making might become important. As protein condensation can be synthetically tuned^108^, our findings suggest a route to controlling such decisions.

## Limitations

There are currently no known mutations that specifically disrupt P-body condensation without affecting protein structure/function. The limitation is common in condensate biology and creates two problems. First, the functional effect of a mutation might be due to it affecting the core function of a protein rather than condensation of the protein, which could be incidental. Second, it is difficult to delineate the contributions of the mesoscale condensate versus smaller underlying molecular assemblies.

To address these limitations, we took advantage of the redundancy in P-body formation and used multiple mutants to correlate the extent of P-body formation with the extent of the phenotypic effects. While our experiments do not precisely differentiate between condensation of ribonucleoproteins (RNPs) into clusters of a few RNPs and mesoscopic P-bodies visible by microscopy^109^, microparticle tracking experiments suggest that the mutations we used primarily target interactions required for mesoscopic P-bodies and leave sub microscopic assemblies intact^110^. Furthermore, the mutations do not prevent P-bodies from forming when cells are in stationary phase, and overexpression of other proteins can restore normal P-bodies in other conditions, indicating that they contribute to a redundant assembly network^110^. Further supporting the notion that P-bodies themselves drive the processes we describe here, when senescent mother cells lose P-bodies to their daughters it revives their division rate (**Figure S4H**). This is consistent with daughter cells slowing their division rate or failing to divide after inheriting a P-body^54^. Nevertheless, future structural studies that separate intermolecular interactions that specifically drive condensation from those that affect the function of the protein in the dilute phase will help to clarify the different contributions.

We do not yet know how Whi3 condensation promotes *CLN3* inhibition. Based on our genetic analyses, we favour a model where Whi3 and *CLN3* are bound in the condensate, as discussed above (**Figure 7E**). However, in this study we did not visualise RNA, so other interpretations are possible, although the qualitative aspects of the model are independent of them. Another question raised by our findings is, what causes age-dependent P-bodies to form in the first place? They require Asc1 to form **(Figure 5G**), but further work will be needed to understand the molecular basis. Furthermore, P-bodies must have roles in lifespan control separate from their role in decision making, as they are required for a normal lifespan (compare **Figures S5C and S5D**).

## Methods

Further information and requests for resources and reagents should be directed to the lead contact, Yves Barral.

### Experimental model

*S. cerevisiae* strains used in this study were constructed in the BY4741 background, which is isogenic to S288C^111^ (https://www.yeastgenome.org/strain/by4741).

## Method details

### Yeast strain construction

Fluorescent protein tagging, gene deletions and truncations were done at the endogenous genetic loci using classical genetic approaches^112^. Genetic modifications were confirmed using two PCR reactions overlapping each junction. CRISPR-Cas9, with pML104^113^, was used to generate the Whi3-ΔQ strain for the Cln3 expression reporter experiments. A repair template that removed the part of the WHI3 coding sequence that encodes: QQQSQQQQQQPQQPQQHSTQKHSPQQCNQQQVN was used. A yeast strain containing the Cln3 reporter was a kind gift from Matthias Heinemann. The reporter was amplified from genomic DNA of the original strain (YSBN6-CLN3-P2A-sfGFP-KANMX), and integrated into wild type cells or Whi3-ΔQ mutant cells. Whi3-ΔQ-3sfGFP and Whi3-ΔQN-3sfGFP were generated in a previous study^15^. In these cases, the prion-like domains were replaced with 3xHA tags^15^. pHR-mCh-Cry2olig was a gift from Clifford Brangwynne (Addgene plasmid #101222; http://n2t.net/addgene:101222; RRID: Addgene_101222) and was used as a PCR template to generate an mCh-Cry2olig fragment that was inserted into AscI-cut (R0558S, NEB; Ipswich, MA, USA) pYB968, to generate the optoYeast plasmid. This was used as a PCR template for tagging Whi3 and other P-body associated proteins. To generate a control strain expressing the mCh-Cry2olig tag alone, the tag was cloned into pRS303^114^ upstream of a WHI3 promoter, using Gibson Assembly^115^ with a Gibson Assembly Master Mix (E2611L, NEB; Ipswich, MA, USA). This plasmid (optoYeast_control) was linearised with BstXI (R0113S NEB; Ipswich, MA, USA) and transformed into BY4741 yeast. Cloning PCR reactions were with iProof polymerase (#1725301 BioRad; Hercules, CA, USA) and confirmatory reactions were with Taq polymerase (M0273E, NEB; Ipswich, MA, USA). All standard oligos were ordered from Microsynth (Microsynth AG; Balgach, Switzerland). Yeast strains were stored at -80°C in yeast extract peptone dextrose (YPD) and 17.5% glycerol.

### CLN3 allele replacement

We generated an endogenous *CLN3* mutant with reduced Whi3 association but that is translated to give native Cln3 protein^36^. CRISPR-Cas9 introduced a double stranded break in the CLN3 locus, which was repaired with a hygromycin resistance cassette with flanking CLN3 5’- and 3’-UTRs to generate a *cln3Δ* strain that retained the CLN3 5’- and 3’-UTRs. CRISPR-Cas9 was used to cut the hygromycin cassette (hphNT1) and a synthesised CLN3 allele (Genscript; Piscataway, NJ, USA) with 14 GCAT replacements was used to repair the cut by replacing hphNT1 through homologous recombination. In the 5’-UTR, 6 GCAT sites were mutated to CGTA. In the coding sequence the 6 mutations, in order of 5’ to 3’ appearance, were: GCAT to GCAC (MH to MH); GCATTA to GCCCTT (AL to AL), GCATT to GTATC (SI to SI), GCAT to GCTT (AS to AS), GCAT to GTAT (SI to SI), GCAT to GTAT (SI to SI). In the 3’-UTR, 2 GCAT sites were mutated to CGTA. We referred to this *CLN3* allele as *CLN3*^mGCAU^ as was done originally for a plasmid version^36^.

### Cell culture

All cultures were grown at 30C with shaking at 180rpm, in synthetic complete medium (SC medium; ForMedium, Norfolk, UK), containing 2% glucose.

### Cell culture – microfluidic ageing experiments

Cells were cultured in SC medium supplemented with 0.1% Albumin Bovine Serum (protease free BSA; Acros Organics, Geel, Belgium). Prior to ageing experiments, yeast strains were taken out of storage and grown on yeast extract peptone dextrose (YPD) plates containing 2% glucose for 24 hours. The plated cells were used to inoculate 10 ml pre-cultures that were grown for 6 hours and used to inoculate a second set of 10ml cultures which were grown for at least 16 hours to an optical density 600 (OD_600_) less than 0.6. The young cells were diluted into fresh growth medium to an OD_600_ of 0.1 and, using a 1 ml syringe, were loaded onto a microfluidic device (see ‘Microfluidic device’ section below) that had been degassed in a desiccator at a pressure of 90 millibars for 1 hour prior to being primed with fresh growth medium. The microfluidic device was continuously flushed with fresh medium at a constant flow rate of 10 μl/min using a Harvard PHD Ultra syringe pump (Harvard Apparatus; Holiston, MA, USA) with up to eight 60 ml syringes (one for each of 8 sample lanes in the device), with inner diameter 26.7 mm (Becton Dickinson; Franklin Lakes, NJ, USA). The syringes were connected to 1/2” LS20 Luer Stubs (Phymep; Paris, France), which were connected to 0.03” inner diameter, 0.09” outer diameter Masterflex Transfer Tygon tubing (MFLX06419-03, Avantor; Radnor Township, PA, USA), which was connected to 0.3 mm inner diameter, 0.76 mm outer diameter PTFE tubing (Fisher Scientific #11919445, Adtech Polymer Engineering, Gloucestershire, UK). The joint between tubes was glued with UHU All Purpose Adhesive Super (UHU GmbH & Co. KG; Bühl, Germany) to prevent leaks. The end of the thin tubing was cut diagonally to help insert it into the inlet holes of the microfluidic device.

### Microfluidic device – design and fabrication

We developed a microfluidic device designed to trap yeast cells and monitor the ageing process through continuous media flushing. The device’s design draws inspiration from previous publications on microfluidics dissection ^29,40,116^, and the design drawing file is available upon request.

The device was constructed using soft lithography, comprising a cover glass and PDMS (Polydimethylsiloxane, Sylgard 184, Dow Corning), replicated from SU-8 fabricated wafers through replica molding. The wafer, featuring microstructures replicated via photolithography using negative photoresists (SU-8, Microchem Corp.), produced an array of cell traps with SU-8 2005 (height = 5 µm), while the inlet and outlet were formed using SU-8 3025 (height = 30 µm). Following wafer fabrication, surfaces underwent silanization to enhance PDMS detachment during replica molding. Silanization involved placing the wafers in a desiccator with 100 µl 1H,1H,2H,2H-Perfluorodecyltrichlorosilane (ABCR GmbH), incubating for 24 hours in a vacuumed environment. For PDMS replication, the PDMS base and curing agent were thoroughly mixed in a 1:10 w/w ratio, degassed in a vacuum chamber to eliminate air bubbles, poured onto the SU-8 master mold, and cured in an oven (80°C for at least 2 hrs). The cured PDMS was then delicately peeled off the mold, and air holes were punched using a needle. PDMS and cover glass surfaces were activated by plasma treatment. Immediately following activation, the PDMS and cover glass were aligned and bonded to form the device. For some initial exploratory studies (Figures S4A-G) an older design was used^116^.

### Timelapse microscopy

The microfluidic device was securely mounted on the stage of an inverted Nikon Eclipse TiE microscope (Nikon Instruments; Tokyo, Japan), incorporating a hardware-based automated focusing system, specifically the Perfect Focus System (PFS). Placed within a controlled incubation chamber (Life Imaging Services; Basel, Switzerland) at 30°C, imaging was conducted utilizing a Plan Apo 60 × 1.4 NA oil immersion objective (Nikon Instruments; Tokyo, Japan). Micro-manager open-source software^117^, managed the application of the requisite excitation and emission filters and captured multiple fields of view per time point with a motorized XY-stage. Depending on the experiment, the light source for fluorescence imaging was a Spectra-X LED Light Engine (Lumencor; Beaverton, OR, USA) or a CoolLED pE-300 (CoolLED; Andover, UK).

We acquired brightfield images every 15 minutes in a single focal plane. In general, a stack of 7 fluorescence images with 0.5 μm spacing was acquired at 2 hours then every 12 hours if the experiment involved imaging fluorescent proteins. For optogenetics experiments the imaging sequence was different (see ‘Optogenetics’ section).

### Optogenetics

Optogenetic ageing experiments were done on a single microscope with a CoolLED pE-300^white^ light source (CoolLED; Andover, UK), described in the ‘Timelapse microscopy’ section. Brightfield images were acquired every 15 minutes in a single focal plane. Every 1 hour 15 minutes an mCherry fluorescence image was acquired in a single central focal plane, then cells were irradiated for 1 s with blue 405 nm light at 50% LED power via a DAPI HC filterset (AHF analysentechnik AG, Tübingen, Germany) in the same focal plane.

To image protein condensation dynamics, young cells were loaded onto CellASIC ONIX Y04 microfluidic plates (Merck; Rahway, NJ, USA). The cells were perfused with SC medium containing 2% glucose, using a CellASIC ONIX microfluidic platform (Merck; Rahway, NJ, USA). 3.5 psi of pressure was applied to drive the flow. mCherry fluorescence and brightfield images were acquired before, then every 5 minutes after a 1 s pulse of blue 405 nm light at 50% LED power. To quantify protein condensation, cells were segmented using the YeaZ convolutional neural network model for brightfield images of yeast cells^118^. The dispersion index of the mCherry fluorescence signal was then measured and further analysed in R (see, ‘Quantifying protein condensation’).

To test for possible effects of repeated rounds of optoWhi3 condensation, an ageing experiment was set up where genetically identical optoWhi3 cells were divided into two groups. At the start of the experiment, one group of cells was exposed to a 1 s pulse of blue light to induce optoWhi3 condensation, and the dissolution of optoWhi3 condensates was monitored in that group of cells. The same group of cells was exposed to the blue light pulse again every 1 h 15 min, to repeatedly induce optoWhi3 condensation. After 24 hours, dissolution of optoWhi3 condensates was monitored again in the cells that had been repeatedly exposed. The second group of cells, which were now old, but had never been exposed to blue light, were then exposed to a 1 s pulse of blue light to induce optoWhi3 condensation, and the dissolution of optoWhi3 condensates was monitored. Senescent cells were excluded from the analysis to avoid possible confounding effects due to endogenous age-induced optoWhi3 puncta.

### Optogenetic plating experiment

Cultures were grown overnight in SC medium (SC medium; ForMedium, Norfolk, UK), 2% glucose at 30°C to an OD_600_ less than 0.6, then diluted to an OD_600_ of 0.004 in fresh medium. 300 μl of each diluted culture was spread on SC plates (SC medium; ForMedium, Norfolk, UK), containing 2% glucose, using glass beads. A blue 5 mm (T-1 3/4) LED with a 465 nm dominant wavelength (L-7113QBC-D, Kingbright; City of Industry, CA, USA) was held above plates, using a custom-made acrylic lid that held the LED 7 mm above the surface of the centre of the plate. The LED was controlled with an Arduino UNO Rev3 microcontroller (Arduino; Monza, Lombardia, Italy), using Arduino IDE software version 1.8.15, and a basic sketch based on code in the public domain: https://www.arduino.cc/en/Tutorial/BuiltInExamples/Blink. The LED was turned on every 3 seconds for 3 seconds, which was found to be a power where wild type cells in the small most intensely irradiated region of the light path had reduced growth. The whole system was placed inside an ECHOtherm benchtop incubator (Torrey Pines Scientific; Carlsbad, CA, USA) at 30°C, with a tray of sterile water to maintain humidity. Cells were allowed to grow for 3 days, after which the plates were imaged with a desktop scanner, and samples of cells were scraped from different regions of the plate, resuspended in 2.5 μl of growth medium and wet-mounted on a No. 1, 18 x 18 mm glass coverslip (VWR; Radnor, PA, USA) for immediate fluorescence microscopy on one of the microscopes described in the ‘timelapse microscopy’ section.

### Cln3 production assay

We bleached the sfGFP reporter with 20x 600ms pulses 488nm light with 3 s intervals between them and monitored recovery of the GFP signal every 15 minutes. The bleaching sequence did not affect the lifespan of the cells. We performed the experiment on young (3 h after loading) and old cells (48 h after loading). Background subtraction based on a field of view that did not contain cells accounted for possible intensity fluctuations due to the microscope. For each field of view, there was a specific delay time between bleaching and recovery monitoring, which ranged from 0 s to 162 s. These times were accounted for in the analysis. To average the recovery of multiple cells, the intensity (I) during recovery was normalised against the intensity after bleaching (I_0_):

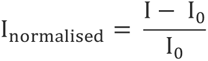

To get the initial rate of Cln3 production, a linear model was fit to the first three data points.

### Quantifying Whi3 enrichment in P-bodies

Age-induced P-bodies were manually segmented in Fiji^119^ from the fluorescence channels of ageing movies using only Edc3-mCherry as a marker of P-bodies. Pixel scaling was fixed with a range that was suitable for viewing the P-bodies, to ensure consistent segmentation. This generated a set of ROIs corresponding to P-bodies, ‘P-body ROI’. Each ROI was moved into the cytoplasm to an area without a P-body and used to generate a second set of ROIs, ‘cytoplasm ROI 1’. The process was repeated in a new area of cytoplasm without a P-body to generate a third set of ROIs, ‘cytoplasm ROI 2’. The ROIs were linked to the IDs of the individual cells throughout. We measured the mean fluorescence intensities in these ROI sets in the Edc3-mCherry and Whi3-3sfGFP channels. We quantified Whi3 enrichment in P-bodies by dividing Whi3-3sfGFP intensity in the P-body ROI by Whi3-3sfGFP intensity in the cytoplasm 1 ROI, on a single cell basis. As an internal control we repeated this between cytoplasm 1 ROIs and cytoplasm 2 ROIs.

### Pheromone switching experiments

We optimised a simple method to switch cells on the microfluidic device to pheromone containing growth medium. For each sample/channel on the device, the outlets of two syringes (the setup for the syringes is described in the ‘Cell culture – ageing experiments’ section) were connected to a 3-way stopcock (#19406-49, Cole-Parmer; Vernon Hills, IL, USA). The syringes were connected to the stopcock with 1/16” barb Luer locks (#45513-00, Cole-Parmer; Vernon Hills, IL, USA) and (#MFLX45508-01, MasterFlex Avantor; Vernon Hills, IL, USA). One syringe pump was turned on, with the flow rate set to 20 μl per minute and with the stopcock set to exclusively allow the syringe to supply the microfluidic device. The second syringe pump was turned off, and the medium in the syringe was blocked from entering the device by the stopcock. To switch the medium, the flow rates of the two syringes were reversed and the stopcock valve was rotated to the allow the new ‘on’ syringe to supply the microfluidic device. After the switch, the flow rate was temporarily increased to 200 μl per minute for 10 minutes, then returned to 20 μl per minute. This allowed the media to be exchanged within 10 minutes, much faster than the pheromone response time of the cells, which happens on a scale of hours. Switching was monitored using an Alexa Fluor™ 680, 3000 MW fluorescent dextran (D34681, Invitrogen; Waltham, MA, USA). The slightly higher basal flow rate of 20 μl per minute in these experiments was used to prevent cells from clumping due to changes in their morphologies after shmooing.

### Quantifying protein condensation

We quantified the dispersion index of pixel intensities in individual cells (the ratio of the variance to the mean) as a readout for protein condensation.

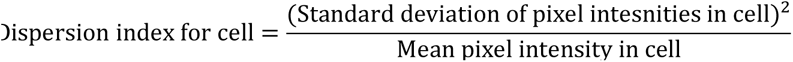

Where comparisons were made across experiments, the dispersion index was normalised to an internal standard: cells from the same ageing experiment with a wild type background except for the tagged protein. Pixel intensity variance and mean were measured using Fiji and exported to R for analysis. For old cells, we measured dispersion index in the last 5 divisions. If multiple fluorescence images were acquired in the last 5 divisions, the image corresponding to the youngest age in those last 5 divisions was used. For young cells, dispersion index was measured in the first 0-3 divisions.

### Quantifying Whi3 puncta

The nebulous appearance of endogenous Whi3 puncta was not suited to the dispersion index method because it required the protein of interest to be highly portioned between the dense and dilute phases. Thus, Whi3 puncta were scored by eye from background subtracted images (Fiji, 200 pixel rolling ball radius). Regions of intense Whi3 fluorescence had to span at least 2 focal planes with a 0.5 μm separation to be considered puncta.

### Division time analysis

Division times were annotated manually from ageing experiment movies, as the time between the visible start of budding (usually visible as a dark patch), to cytokinesis. Cytokinesis was defined as the separation of the cytoplasm of the mother from the daughter. In old cells this was often visible as a dark line separating the mother and daughter (probably the secondary septum), after which another division started, even if the original daughter cell was only released later. In younger cells, formation of the septum and release of the daughter cell were indistinguishable. All individual cells in a field of view were systematically cropped out prior to analysis. During the analysis, if a cell happened to be washed out of the trap, it was discarded. Annotated division times were analysed further using R. A quality control step was applied to correct possible annotation mistakes. This involved searching and correcting negative division times and double-checking division times that were very short or very long.

### Cell cycle analysis

We filtered for cells’ last 10 cycles with detectable G1s (we did not detect most G1s in young cells as the mean G1 duration of a young mother cell is close to our sampling interval of 15 minutes^120^). The full cell cycle length as well as G1 and division times were plotted as histograms.

### Senescence entry analysis

First, all divisions were classified as either normal (< 1.5 h) or ‘slow’ (>= 1.5 h). Then, groups of 3 or more consecutive slow divisions were labelled as the senescent state. Next, transitions from the non-senescent to the senescent state were labelled with 1, and transitions from the senescent to the non-senescent state with -1. The cumulative sum of the transitions was converted to a percentage of the number of cells, which was plotted against the division number. The resulting plots show the percentages of cells that are senescent after a given number of divisions. Statistical analysis of the distributions was done using the exact two-sample Kolmogorov-Smirnov test.

### Lifespan/survival analysis

The microfluidic device we developed allowed us to retain > 95% of cells until their death, which is important to prevent biasing data. All cells in a field of view were analysed. Those that were washed away were also incorporated into our analyses. Cell survival was assessed by plotting Kaplan-Meier survival curves^121^ (non-parametric statistic for estimating the survival function from lifespan data). Kaplan-Meier analysis may provide a better estimation of lifespan in yeast ageing experiments because it considers censored data (cells that are randomly washed away)^122^.

R packages survival^123^ and survminer^124^ were used for Kaplan-Meier analysis and generating survival plots. For summaries of percentage lifespan reduction (Figure 6M), error propagation was done using the pairwiseCI (version 0.1-27) R Package^125^, using the MOVER-R method (https://www.rdocumentation.org/packages/pairwiseCI/versions/0.1-27/topics/MOVERR) for dividing confidence intervals^126^.

### Computational disorder prediction

DISOPRED3 https://github.com/psipred/disopred^127^ was used to predict structural disorder from protein amino acid sequences.

### GO term analysis

We used GO Slim Mapper (https://www.yeastgenome.org/goSlimMapper) which is available through the Saacharomyces Genome Database (SGD)^128^. We searched query genes for all Yeast GO-Slim processes.

### Data wrangling and visualisation

ggplot2^129^ was used to create all the plots in this study. Data wrangling prior to plotting was done using the other tidyverse packages^130^. A range of R packages were used to facilitate analysis, statistics, and visualisation. These were: svglite^131^, slider^132^, Rmisc^133^, Hmisc^134^, viridis^135^, ggrepel^136^, ggforce^137^, colorspace^138^, and scales^139^. All analyses and visualisation were done in the RStudio environment^140^. Illustrations were created with Inkscape^141^.

### Quantification and statistics

No statistical methods were used to predetermine the sample sizes. All ageing experiments were repeated at least twice with very similar results. As replicative lifespans can vary by one or two divisions between individual experiments, analyses from individual experiments are shown. Relative differences between wild type and mutant strains were the same between experiments. Details of the statistical methods used are in the experimental methods sections.

## Acknowledgements

We thank Daniel Smith for making the custom acrylic lid used in the optogenetic plating experiments. We thank Laura Mariotti, Vamshidhar Gade, Stephanie Weber and Hugo Stocker for detailed comments on the manuscript, Aliaksandr Damenikan for discussions on decision making, and the Barral and Kroschewski labs for their feedback on the project and manuscript.

## Author contributions

T.R.P. and Y.B. conceptualised the project, designed experiments, interpreted the results, and acquired funding. T.R.P. performed the experiments, analysed and visualised the data, and drafted the manuscript. S.S.L. designed and fabricated the microfluidic chips and provided microscopy support. All authors reviewed and edited the manuscript.

Y.B. acknowledges support from ETH Zürich and from the Swiss National Science Foundation (31003A-105904 project funding grant).

T.R.P. was supported by an ETH Zürich Postdoctoral Fellowship (18-2 FEL-07) co-funded by the Marie Skłodowska-Curie Actions COFUND Program, EMBO Postdoctoral Fellowship (ALTF 776-2018) and Marie Skłodowska-Curie Postdoctoral Fellowship (H2020-MSCA-IF 844867 MemoryAggregates), and the NCCR RNA and Disease Parental Leave Support Measure.

## Declaration of interests

The authors declare that they have no competing interests.

## Notes

### Competing Interest Statement

The authors have declared no competing interest.

