## Supplementary figures and images for "A protein condensation network contextualises cell fate decisions"

### Supplementary Figures

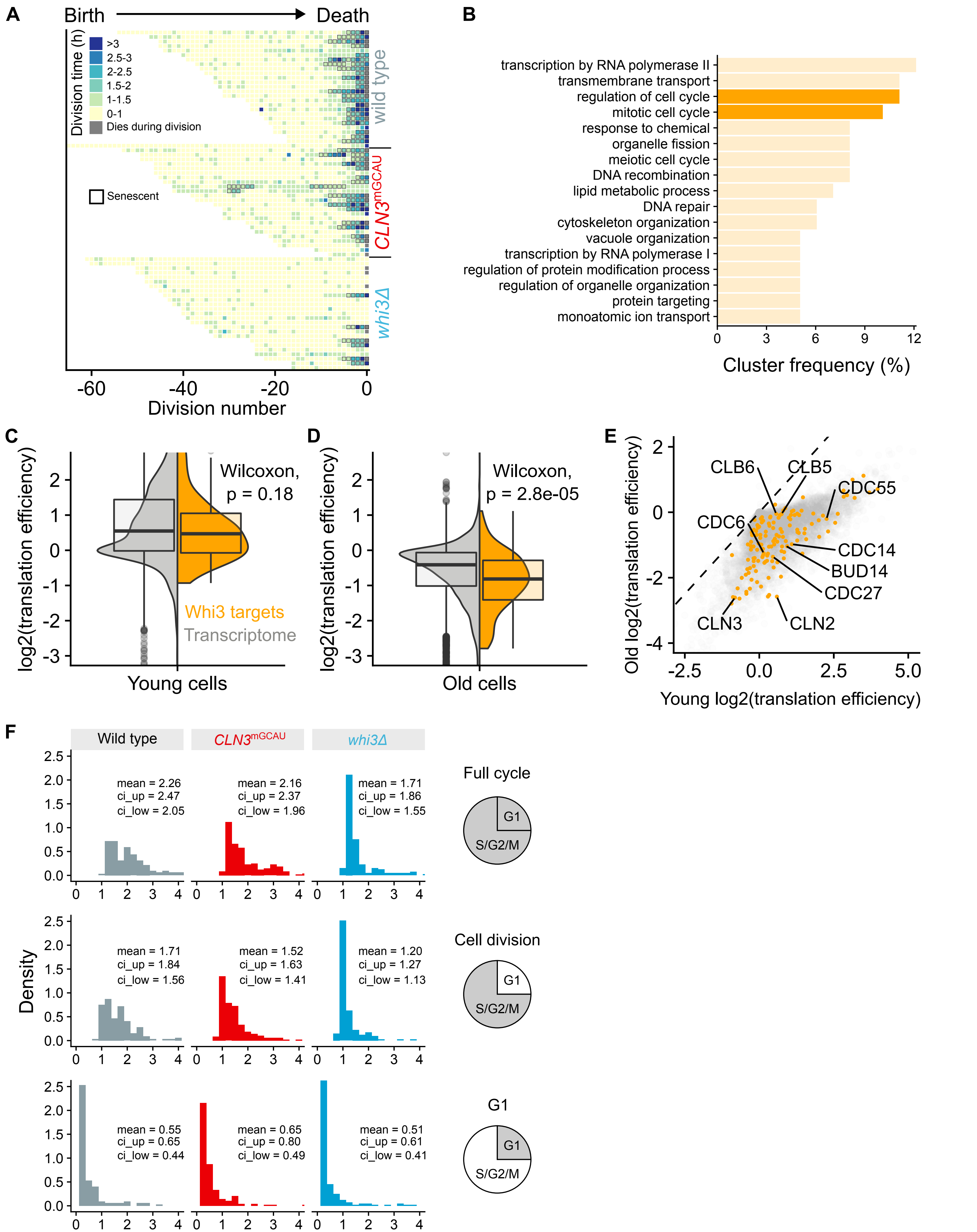

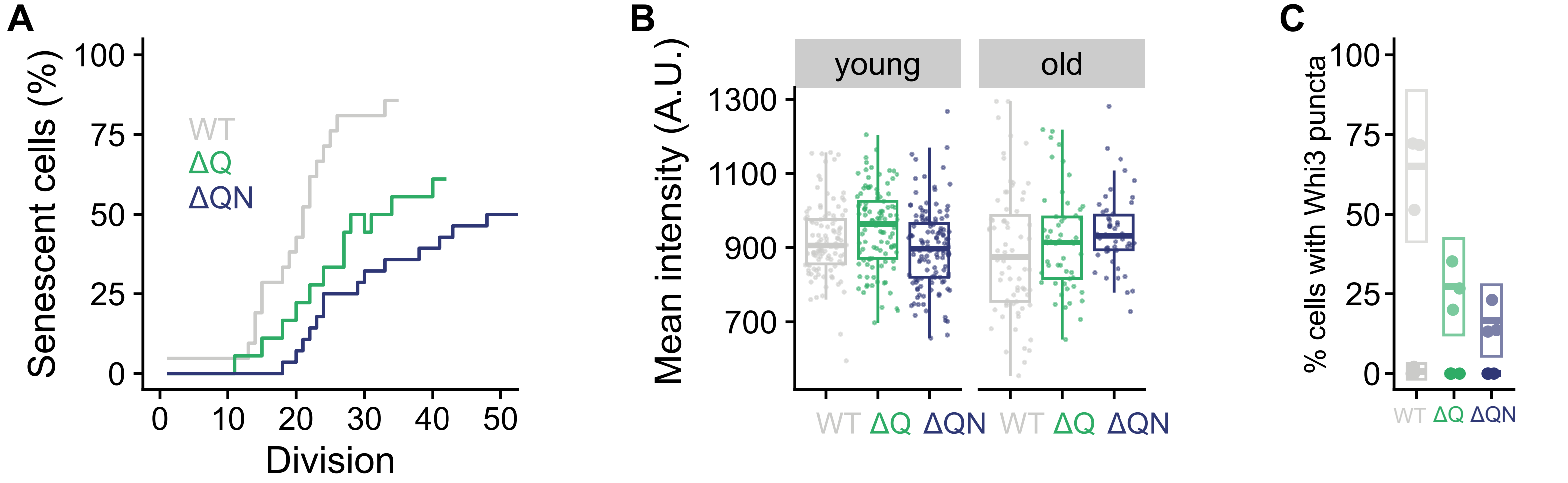

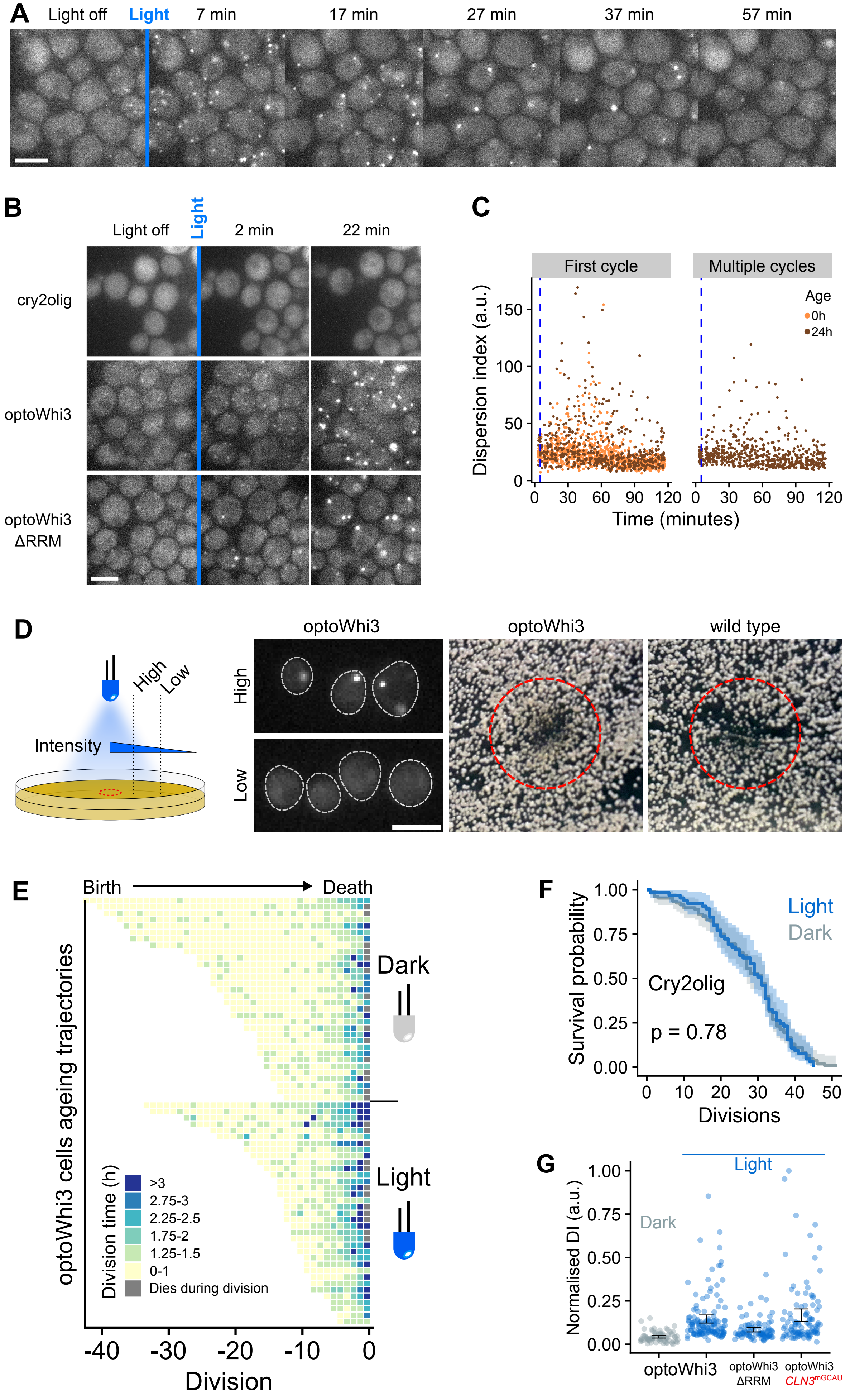

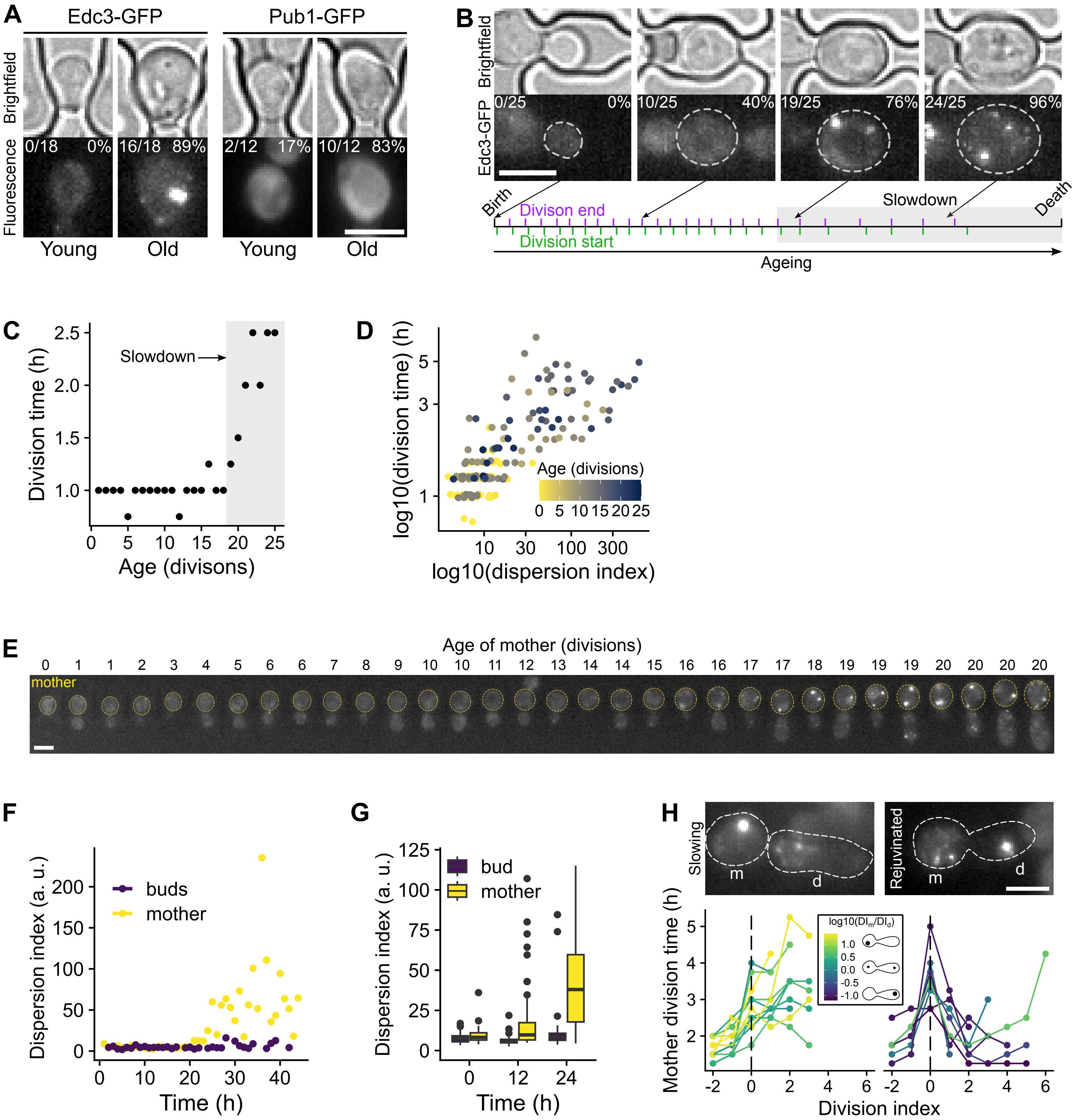

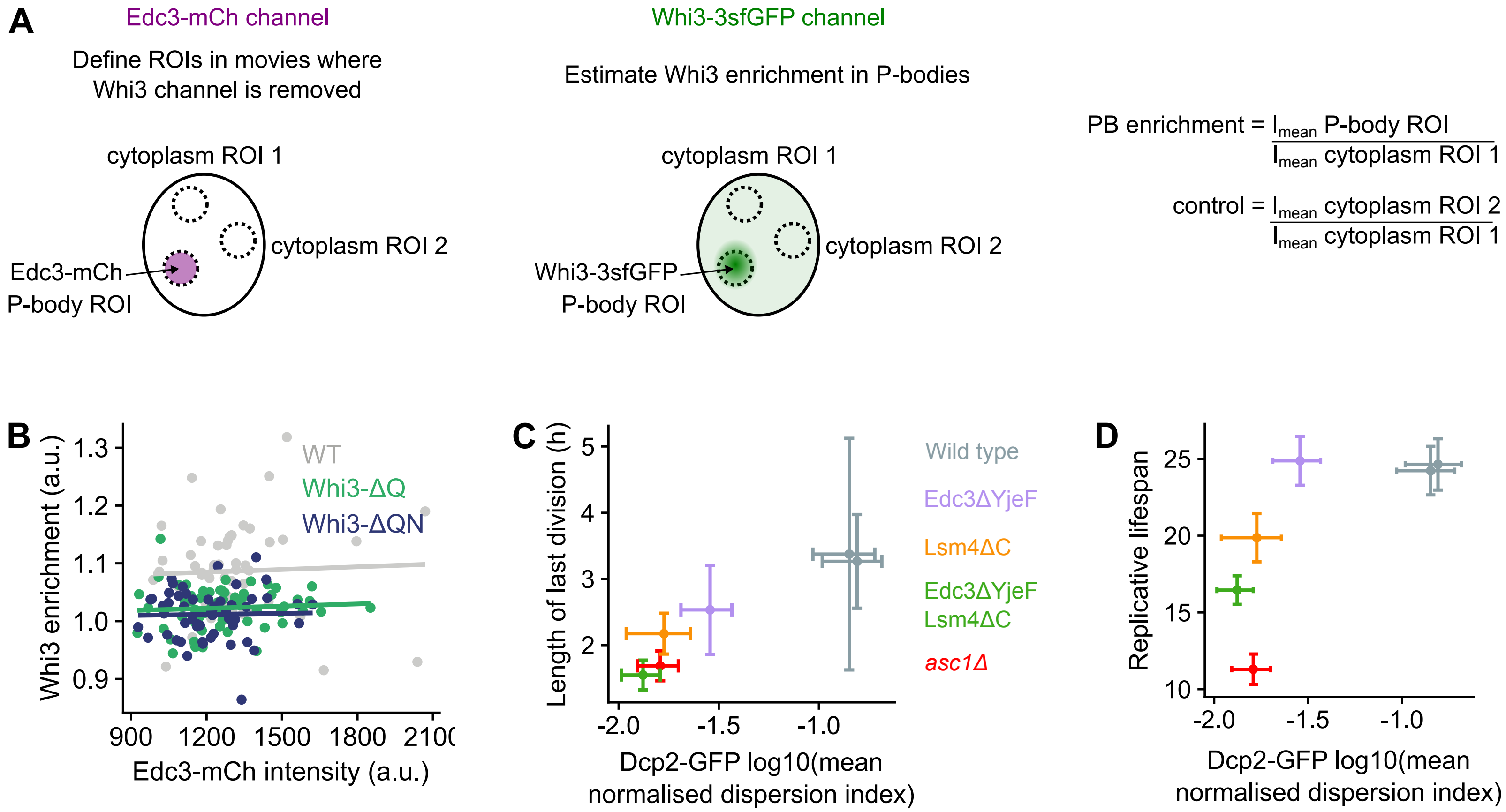

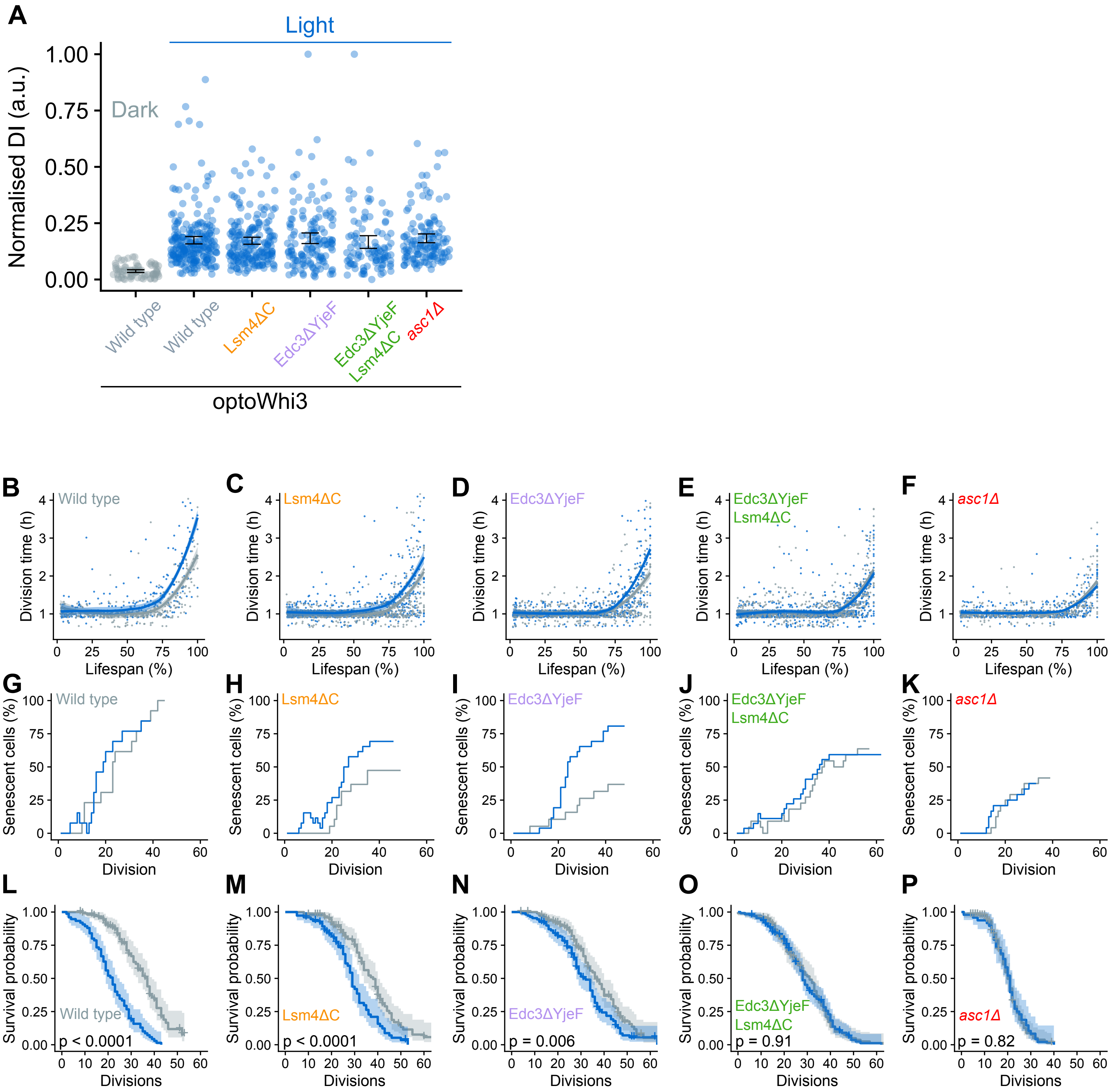

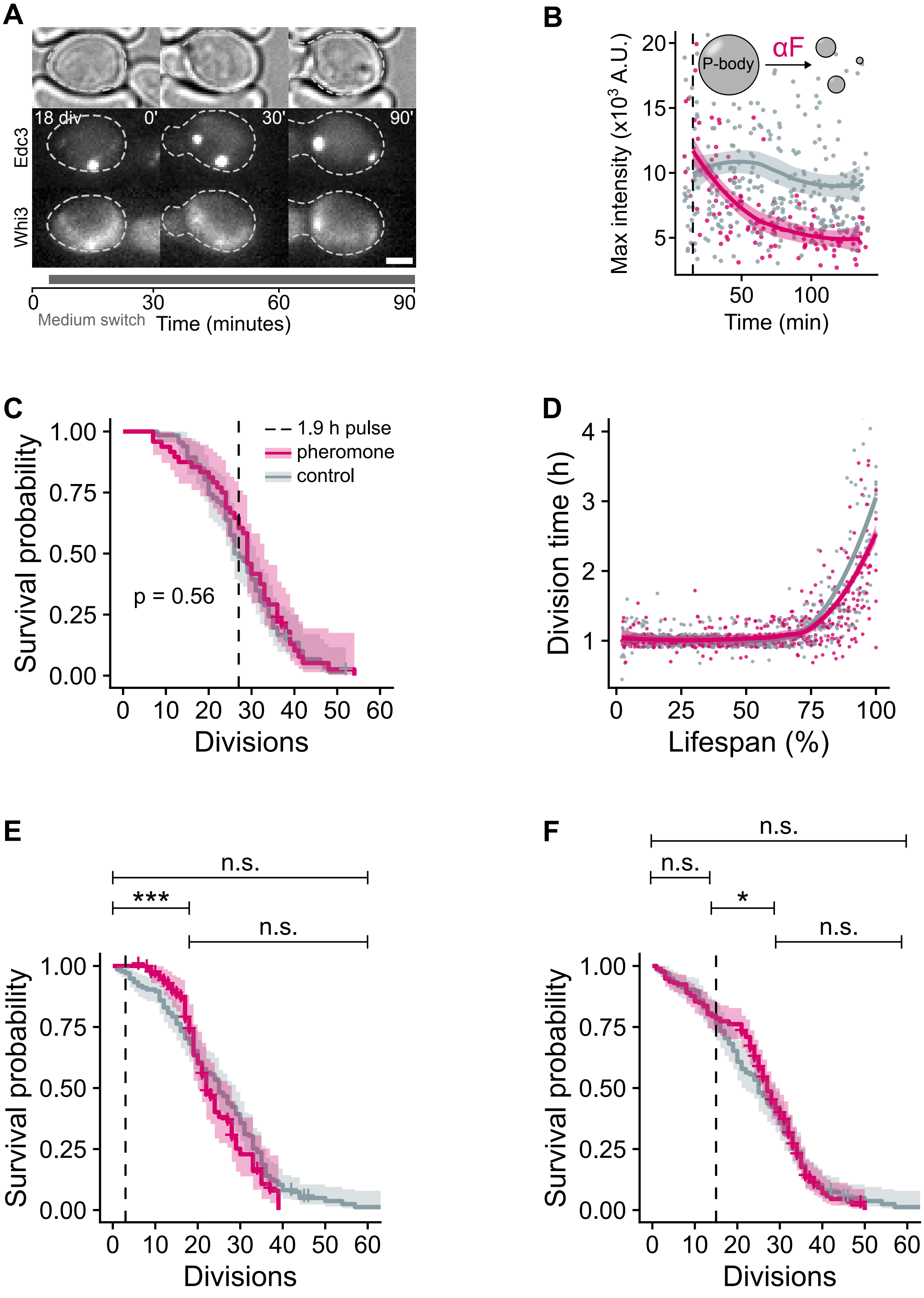
